# Structural insights into DNA annealing and recombination by herpesviral DNA-binding proteins ICP8 and BALF2

**DOI:** 10.1101/2025.06.25.661518

**Authors:** Girish R. Apte, Shuvankar Patra, Mariusz Czarnocki-Cieciura, Aqsa Jabeen, Justyna Jackiewicz, Jarosław Poznański, Marcin Nowotny, Małgorzata Figiel

## Abstract

The DNA-binding proteins of herpesviruses are key factors involved in multiple stages of DNA replication and recombination. The well-studied ICP8 protein of Herpes Simplex Virus 1 binds single-stranded DNA in a cooperative manner and facilitates the annealing of complementary DNA strands in a process that involves the formation of double helical filaments. Moreover, it cooperates with UL12 alkaline nuclease to execute recombination between homologous DNA sequences. In this work, we present the structures of ICP8 and its homolog BALF2 of the Epstein-Barr Virus in complexes with DNA. These structures reveal a conserved conformation of the bound DNA in which several bases are rotated away from the protein, enabling homology search and base-pairing. The crystal structures of ICP8-DNA depict the pairing between two DNA strands, showing the flexibility of base-pairing. In the structure of the double filament of ICP8, we visualize the interactions between protein subunits within and between the antiparallel strands. Finally, we present the first complete structure of a ternary recombinase complex composed of ICP8 and UL12 bound to a model DNA. This structural landscape illustrates the different stages of the DNA recombination process in herpesviruses, shedding light on molecular mechanisms of their replication.

## INTRODUCTION

Replication of the double-stranded genome of herpesviruses requires the concerted action of a set of proteins. Among them, six core replication proteins are essential in all herpesviruses. In Herpes Simplex Virus (HSV-1) these include the DNA-binding protein (ICP8), heterodimeric DNA polymerase (UL30, UL42), and a tripartite helicase-primase complex (UL5, UL8, UL52)^1^. In alphaherpesviruses, this core set additionally includes the origin-binding protein (UL9).

HSV-1 infection causes a reorganization of the host nucleus, which involves the formation of prereplicative sites and, subsequently, replication compartments. Functional ICP8 is instrumental in these processes^2,3^. The formation of prereplicative sites critically relies on its ability to form protein-only double filaments^4^. Moreover, ICP8 serves as a multi-purpose cofactor in DNA replication, as it interacts physically or functionally with all key components of the DNA replication machinery. It cooperates with UL9 to open the origin of replication^5,6^. Through its interaction with the UL8 component, ICP8 stimulates the activity of the helicase-primase complex^7,8^ and cooperates with it to initiate the formation of prereplicative sites^9^. Finally, DNA synthesis by the HSV-1 polymerase is stimulated in the presence of ICP8^10,11^.

ICP8 binds to single-stranded DNA (ssDNA) with an estimated footprint of 10-15 nt and holds it in an extended conformation^12–14^. The DNA binding is cooperative leading to protein-DNA filament formation^14^. ICP8 can both destabilize short double-stranded DNA (dsDNA) helices^15^ and mediate strand annealing^16–18^. This annealing involves interaction between ICP8-coated DNA strands and its intermediates can be visualized as supercoiled structures^19^. ICP8 can promote strand transfer between DNA molecules^16^ and a ssDNA fragment coated by ICP8 is capable of invading dsDNA^20^.

Like many dsDNA viruses, HSV-1 uses recombination-dependent replication (RDR)^21^ which enables genome isomerization and offers a way to restart replication after replication fork collapse^22^. The branched intermediates of HSV-1 RDR are likely produced through the combined action of ICP8 and the 5′→3′ alkaline nuclease, UL12^23^. ICP8 and UL12 physically interact^24^ and can jointly mediate a strand exchange reaction between a linear dsDNA and circular ssDNA in vitro^25^. ICP8 increases the processivity of UL12 and specifically stimulates its nuclease activity on dsDNA^26^. This functional association of ICP8 and UL12 is reminiscent of the two-component recombinase systems found in other dsDNA viruses, for example, the λRed system of bacteriophage λ and the RecET system of the *E. coli* prophage Rac^25,27^.

Structurally, most of the ICP8 sequence forms a large globular part, which is followed by a flexible linker, a small C-terminal domain, and an unstructured fragment^28^. The C-terminal domain binds to a concave surface of another ICP8 molecule mediating head-to-tail interactions between ICP8 protomers that enable cooperative DNA binding and filament formation. The C-terminal unstructured fragment of approximately 60 residues contains a nuclear localization signal and is a critical element for self-oligomerization of ICP8 into protein-only filaments and for cooperative binding of ssDNA^4,29,30^.

Relatively few reports exist about the ICP8 homolog from Epstein-Barr Virus (EBV), BALF2. Similarly to ICP8, BALF2 contributes to DNA replication at multiple stages of the process^31–33^. Strand displacement activity of BALF2 has been observed on short stretches of duplex DNA^34^. C-terminally truncated version of BALF2 forms dimers at higher concentrations and coats ssDNA in a weakly cooperative manner^13^. Moreover, studies of a BALF2 homolog from another gammaherpesvirus, Kaposi-Sarcoma Herpesvirus (KSHV), ORF6, revealed that it can oligomerize into protein-only filaments capable of binding ssDNA^35^.

The structure of HSV-1 ICP8 has been determined via X-ray crystallography, and a low-resolution model of the double protein-only filament has been obtained using cryo-electron microscopy (cryo-EM). However, no high-resolution structural data are available to visualize the complexes of DNA-binding protein with DNA, with the alkaline nuclease, or the protein-protein interactions within a double-helical filament. This study aims to address this gap through structural characterization of HSV-1 ICP8 and EBV BALF2 in complexes with ssDNA, elucidating their DNA binding mechanisms, homology search processes, cooperation with UL12 to drive recombination, and the details of the interaction between ICP8 subunits within double filaments.

## RESULTS

### The structure of BALF2-DNA complex

Our aim was to elucidate the structural basis of the interactions between herpesviral DNA-binding proteins and nucleic acids to understand their mechanisms of action. We expressed two homologs, EBV BALF2 and HSV-1 ICP8, in insect cells and purified them. To define the structural basis of DNA binding by BALF2, we pursued its cryo-EM structure determination in complex with ssDNA. To capture complexes formed when multiple BALF2 protomers bind cooperatively to a single DNA molecule, we utilized DNA oligonucleotides of varying lengths: 36, 38, 48, 60, 90, and 131 nucleotides. Particles of good quality were observed only with the 60-mer and 90-mer DNA, and the latter was selected for further processing. The particles exhibited a “beads-on-a-string morphology” and the appearance of the 2D class averages indicated considerable flexibility in the interactions between protein subunits bound to individual DNAs (Fig. 1a, Supplementary Figure S1). Therefore, subsequent processing focused on individual protomers interacting with ssDNA, resulting in a BALF2-DNA complex structure at 3.2 Å resolution (Fig. 1b, Supplementary Figure S2). The structure of BALF2 is very similar to the previously determined structure of HSV-1 ICP8 lacking DNA^28^ (RMSD of 2.0 Å over 713 C-α atoms). ICP8 comprises two main elements: a large N-terminal domain and a smaller, α-helical C-terminal domain. The latter is not visible in BALF2-DNA structure, likely due to its mobility. Based on previous structural work, the large N-terminal domain can be divided into three subdomains, termed the head, neck, and shoulder.

**Figure 1.**
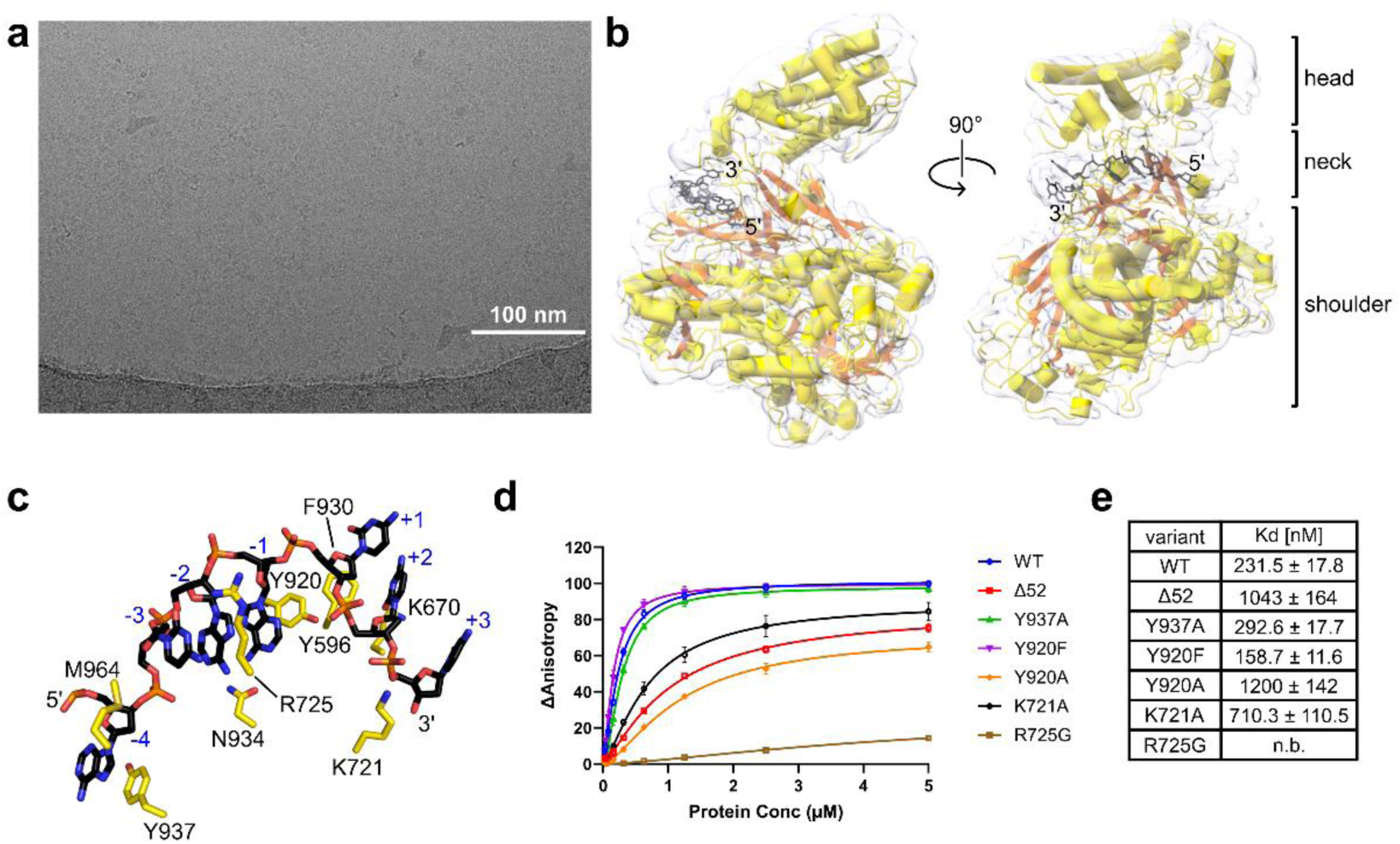
Cryo-EM structure of BALF2-DNA complex and analysis of DNA-binding residues. **a)** Representative cryo-EM micrograph of BALF2 bound to 90-mer ssDNA. Scale bar represents 100 nm. **b)** Reconstructed cryo-EM structure of BALF2 bound to ssDNA. The cryo-EM map shown in gray, with the BALF2 polypeptide chain colored according to its secondary structure elements (helices in yellow, β-strands in orange). DNA is represented as black sticks. **c)** Close-up view of the DNA-binding site of BALF2. Side chains of residues involved in DNA contacts are shown as sticks and labeled. Nucleotides are numbered from −4 to +3. **d)** DNA binding affinities of different BALF2 variants, as determined by fluorescence anisotropy titrations. Measurements were performed in triplicate using a 16-mer ssDNA labeled with Cy5 at its 3′ end. **e)** Dissociation constants (Kds) for the binding of BALF2 variants to the 16-mer ssDNA, calculated from the fluorescence anisotropy data.

We observed density for seven nucleotides accommodated within a positively charged cleft in the neck subdomain (Fig. 1b, 3c). We numbered the positions occupied by these nucleotides from −4 to +3, using negative and positive numbers to define two distinct regions of the DNA. Starting from the 5′ end of the modelled DNA, the nucleobase in position −4 is splayed apart from the base stacking by the sidechain of Met964 and stabilized by an interaction with the aromatic sidechain of Tyr937 (Fig. 1c). The next three bases (positions −3,-2, −1) form stacking interactions and their edges face the protein. The base in position −3 interacts with Arg725, and the base in position −1 with Asn934. The exposed surface of the aromatic ring of the base in position −1 forms interactions with Tyr920. Starting from position+1 the DNA backbone makes a turn with an outward rotation of the base. The sugar ring of this nucleotide is stabilized by an interaction with Phe930. In the subsequent DNA fragment, the edges of the bases are exposed to the solvent and the DNA interacts with the protein through the phosphodiester backbone. The phosphate between positions +1 and +2 is bound by the hydroxyl group of Tyr920; the carbonyl oxygen of Lys670 contacts the deoxyribose in position +2; and the phosphate between positions +2 and +3 interacts with Lys721.

### Identification of key residues responsible for BALF2 binding to DNA

To validate the importance of BALF2 residues participating in DNA binding, we generated protein variants with point substitutions at these positions. We also included a variant lacking the 52 C-terminal residues (Δ52), as deletion of this region in ICP8 has been shown to abolish the cooperative DNA binding. We assessed the affinity of these variants for a 16-mer ssDNA, fluorescently labeled on the 3′ end. Given the length of this oligonucleotide, we expected only one BALF2 protomer to bind. Dissociation constants were estimated from fluorescence anisotropy titration experiments (Fig. 1d, e). The wild type protein bound to the DNA with a K_d_ of 231.5 nM. Substituting Tyr920 with phenylalanine strengthened the binding to DNA (K_d_ of 158.7 nM), whereas replacing it with alanine significantly reduced affinity (K_d_ of 1200 nM). These results highlight the importance of the interactions that stabilize the last nucleotide prior to the rotation in the DNA strand, in particular the base stacking with the aromatic ring of Tyr920. The K721A variant showed reduced DNA affinity (K_d_ of 710.3 nM) likely due to the loss of the lysine sidechain interaction with the phosphate backbone between positions +2 and +3 in the rotated-out region. Substitution of Arg725 with glycine eliminated DNA binding entirely, indicating that stabilizing the portion of the bound DNA upstream of the trajectory change is crucial for efficient binding. Conversely, alanine substitution of Tyr937, which stacks with the nucleotide at position −4 had a minimal effect on DNA binding (K_d_ of 292.6 nM). Finally, DNA binding was strongly reduced in the C-terminally truncated BALF2Δ52 variant (K_d_ of 1043 nM).

In conclusion, our DNA binding assays confirmed the importance of the residues involved in interactions with nucleic acids and demonstrate that Tyr920 and Arg725 play particularly important roles in stabilizing the DNA.

### Crystal structure of ICP8-DNA complex symmetrical dimer

Next, we focused on structural studies of the ICP8-DNA complex. After extensive crystallization trials, we determined the co-crystal structure of ICP8 with a 14-mer ssDNA containing a random sequence. The asymmetric unit of these crystals contained a two-fold symmetric dimer of ICP8 in which each of the subunits binds a DNA and the DNA-binding clefts of the protomers face each other. The protein-protein contacts in the dimer are mediated by the shoulder subdomains (Fig. 2a), but additional contacts are also formed in the head/neck regions, mediated by two mobile elements which are also involved in DNA binding and stabilization: 555-565 and 989-997. The total buried surface area of the dimer interface is 1020.7 Å^2^. The conformation of ICP8 in our complex structure is almost identical to that in the published apo structure of ICP8 (RMSD of 0.41 Å over 854 Cα atoms)^28^.

**Figure 2.**
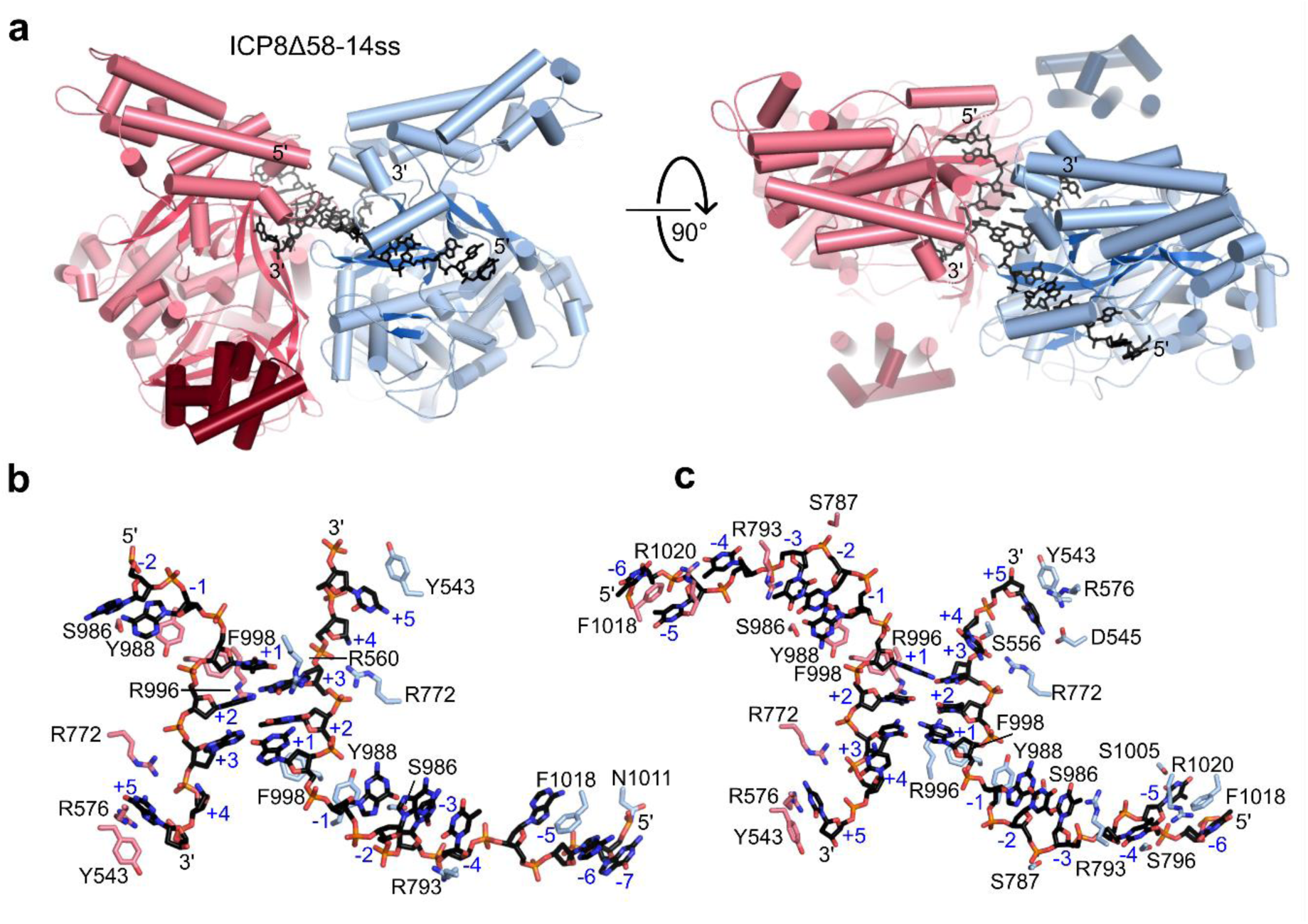
Crystal structure of ICP8-ssDNA complex. **a)** Crystal structure of ICP8Δ58 bound a 14-mer ssDNA. The two ICP8 subunits are colored in shades of pink and blue, respectively, with the C-terminal domains shown in darker shades of the same colors. DNA is represented as black sticks. Rotated view of crystal structure of ICP8Δ58 bound to the 14-mer substrate. **b)** Close-up view of DNA binding in the ICP8Δ58 complex with a random sequence 14-mer ssDNA. Side chains of residues forming contacts with the DNA are shown as sticks, labeled, and colored according to the subunit they belong to. Blue labels denote the numbering of nucleotides relative to the inflection point of the nucleic acid trajectory. **c)** Close-up view of the DNA binding in the ICP8Δ58-8T2C complex. Colors and labels the same as in panel b).

In the ICP8 crystal structure we observed electron density for both the large N-terminal domain and the small helical C-terminal domain. The C-terminal domain has negatively charged patches on its surface and it is accommodated in a positively-charged groove between two helical elements of the shoulder subdomain of a symmetry-related molecule of ICP8. A very similar interaction was observed in the apo structure of ICP8^28^. The asymmetric units of both the apo and ICP8-DNA complex structure comprise protein dimers, but these dimers are defined differently; in the apo structure, the two subunits are in a head-to-tail arrangement, with the C-terminal domain of one subunit interacting with the N-terminal domain of the other. However, when symmetry-related molecules are considered, similar interfaces form in both structures. Comparison of the two-fold symmetrical dimers reveals that, in the presence of DNA, the interaction between the protomers becomes tighter, with the head subdomains moving closer together by approximately 13 Å.

Twelve and seven nucleotides could be built into the electron density adjacent to protein chains A and B, respectively. For the analysis of the ICP8-DNA interaction we employed the same numbering scheme, with positive and negative numbers indicating distinct regions of the DNA. Starting from the 5′ end of the visible DNA, the first contacts are made with the non-bridging oxygens of the phosphate groups on either side of the nucleotide in position −7, and are mediated by the backbone atoms of Gly1008 and the sidechain of Asp1011. The aromatic sidechain of Phe1018 stacks with the nucleobase in position −6 and pushes the nucleobase in position −5 out of base stacking with its neighbors (Fig. 2b). The guanidinium group of Arg793 interacts with the deoxyribose in position −4. Tyr988 stacks with the base in position −1, the location where the DNA backbone undergoes a turn, resulting in the outward rotation of downstream bases to expose their edges to the other ICP8-DNA complex. This portion of the DNA strand is primarily bound by the protein via the phosphodiester backbone: the sugar ring at position +1 interacts with Phe998; and Arg772 contacts the phosphate groups between positions +2 and +3 and between +3 and +4. The base at position +5 is stabilized by a stacking interaction with the sidechain of Y543 and forms a contact with the sidechain of Arg576. Due to poor electron density, the bases in positions +4 could not be visualized.

The arrangement of the dimer places the exposed edges of the bases of three nucleotides (positions +1 to +3) of the DNAs in close proximity of the edges of the bases of the corresponding nucleotides bound by the other ICP8 protomer. The two DNA strands in the dimer have antiparallel polarity. As a result, the nucleotides GTG from one DNA strand are placed opposite the nucleotides TCA from the other strand. Nucleotides T(+2) and G(+3) of the first strand base pair with the A(+3) and C(+2) of the opposing strand with a register shift of one nucleotide (Fig. 2b). This leaves the nucleotides in position +1 unpaired, with their bases positioned close to the sidechains of Met991 from both protomers.

We attributed the imperfect base-pairing of the two strands to the random sequence of the DNA oligonucleotide used, so we performed crystallization trials with 14-mer oligonucleotides that contained a short palindromic region of complementarity (GATC) flanked by homopolymeric tracts. In the crystal structure solved for ICP8-8T2C eleven nucleotides of each DNA chain could be built into electron density (Fig. 2c). Many of the interactions between the DNA strand and ICP8 are similar to those described for the structure with random sequence oligonucleotide, however several unique contacts are also observed, mainly in the −5 to −1 region. The base in position −5 interacts with the sidechain of Ser1005; the base of in position-4 makes contacts with the sidechains of Ser796 and Arg1020; and the guanidinium group of Arg793 replaces the base in position −4 as a stacking partner for the base in position −3. The sidechain of Ser787 contacts the phosphate group between positions −3 and −2. Finally, the base in position +5 interacts with the sidechains of Asp545 and Arg576 and stacks with the aromatic sidechain of Tyr543.

In the co-crystal structure with 8T2C DNA, four bases are rotated, and three of them form pairs with opposing bases (Fig. 2c). However, both ICP8 protomers bind the DNA with the same register, precluding proper GATC palindrome pairing and resulting in mismatches: A-C, T-T and C-A. The C-A base pairs adopt Hoogsteen configuration. Identical imperfect base-pairing was observed in the co-crystal structure of ICP8 with the T73C DNA (not deposited), in which the four-base palindrome was shifted by one nucleotide toward the 5′ end. The pattern of base-pairing in the structures with 8T2C and T73C DNA differs from that observed with the random sequence substrate, where nucleotides +2 pair with nucleotides +3, and nucleotide +1 lacks a pair. Intriguingly, the nucleotide preceding the outward rotation is consistently a purine. This suggests that the presence of a purine base in position −1 influences the register in which ICP8 binds the oligonucleotide. This notion is supported by the observation that ICP8 binds the GATC palindrome in the same manner in the 8T2C and 7T3C oligonucleotides. Furthermore, in the structure with the random sequence DNA, one ICP8 copy binds to the DNA strand close to its 5′ end at the cost of not forming the additional interactions with the neck domain.

In all the ICP8-DNA structures, a loop made of residues 989-997 stabilizes the triplet of nucleotides +1 to +3. This loop, visualized only in the ICP8-DNA co-crystal structure, sandwiches the three nucleotide region whose bases protrude outwards. In the structure with the random sequence DNA the sidechain of Met991 is positioned close to the first rotated base of one DNA strand, while its backbone nitrogen contacts the DNA backbone just upstream of the trajectory change. Additionally, contacts between protein and DNA from different protomers are formed by the sidechain of Arg996 from chain B, which interacts with the third rotated base of chain C, and the sidechain of Arg560 of chain A which interacts with the base in position +1 in chain D. In the T8C2 structure, on the other hand, the sidechains of Arg996 from each ICP8 subunit contact only the DNA strands that is bound by this protomer.

In summary, the structures of ICP8-DNA complexes revealed the mechanism of DNA binding and how the protein promotes flexible base pairing between two DNA strands.

### Comparison of the DNA binding by BALF2 and ICP8

The structures of BALF2 and ICP8 in complex with DNA are overall very similar with an RMSD of 1.95 Å over 676 Cα atoms (Fig. 3a-d). The C-terminal domain of BALF2 could not be visualized in the reconstruction, so the way this domain interacts with the N-terminal domain of another subunit between HSV-1 and BALF2 could not be compared.

**Figure 3.**
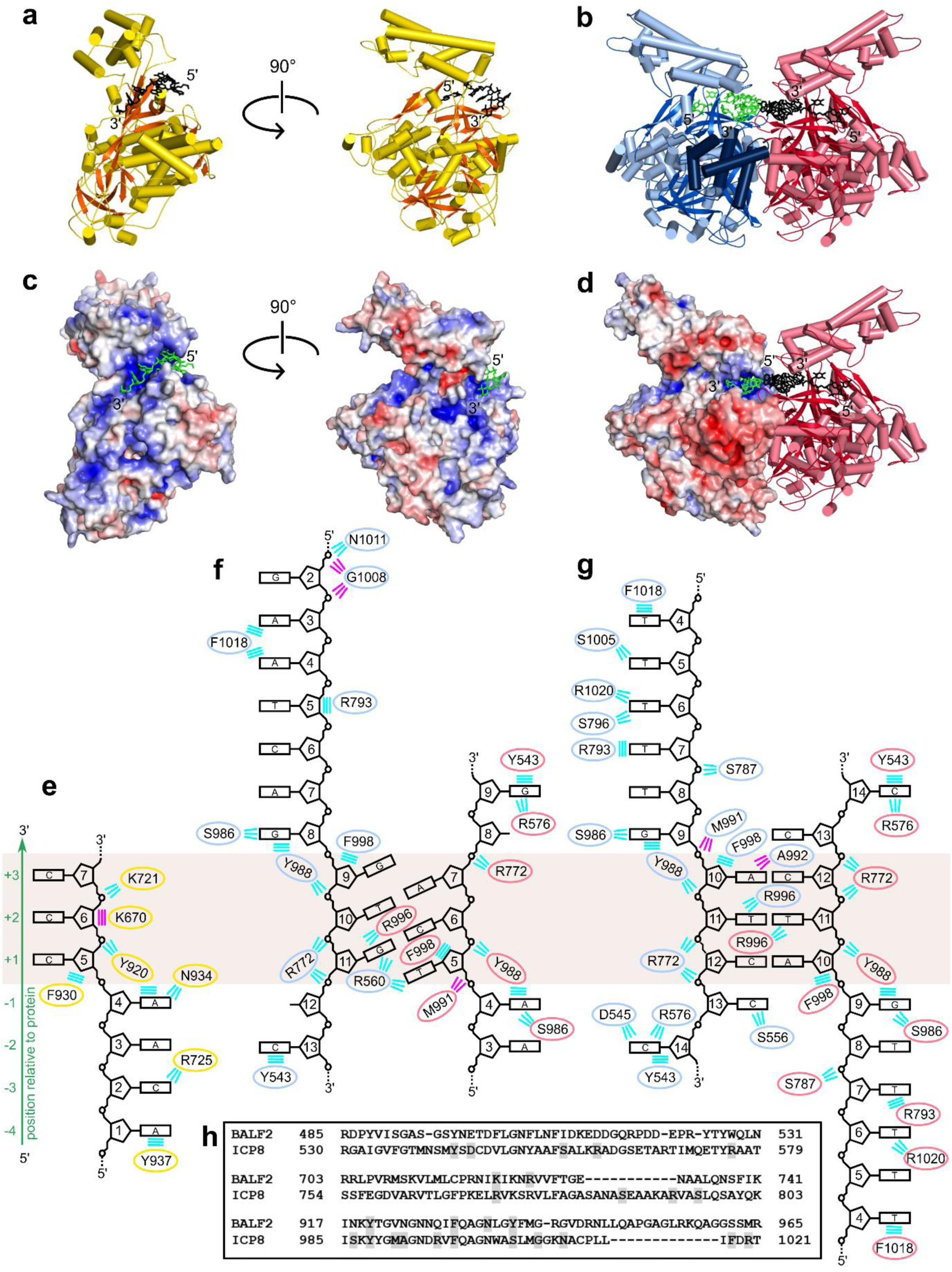
Comparison of ICP8 and BALF2 complexes with DNA. **a)** and **b)** Cartoon representations of the BALF2-DNA and ICP8Δ58-DNA structures, respectively, colored as in Figs. 1 and 2. **c)** and **d)** The BALF2-DNA and ICP8Δ58-DNA structures with one protein chain shown in surface representation, colored according to the calculated electrostatic potential. Bound DNA is shown in stick representation and colored green and black. **e-g)** Protein-DNA interaction schemes for **e)** BALF2-DNA, **f)** ICP8Δ58-14ss, and **g)** ICP8Δ58-8T2C. Nucleotides are numbered either for the modelled region of the DNA (BALF2) or the entire sequence of the oligonucleotide used for crystallization (14ss and 8T2C ssDNAs). Additional numbering for the strand shown with the upward directionality and based on the conformation of the DNA is shown on the left side of the panels (green). The region encompassing positions +1 to +3 is indicated by the beige rectangle. Van der Waals interactions are shown as parallel lines, and polar interactions are shown as radiant lines. Interactions contributed by the sidechains and backbone atoms are shown in cyan and magenta, respectively. Ovals representing sidechains are color-coded according to the protein chain they come from, as in panels a) and b). **h)** Sequence alignment of segments of BALF2 and ICP8 comprising DNA-binding residues (highlighted).

The DNA binding interface is more extensive in ICP8, but several major elements involved in DNA binding are found in both complexes (Fig. 3e-h). The two aromatic residues which stabilize the rotated conformation – Tyr920/Tyr988 and Phe930/Phe998 (from BALF2/ICP8, respectively) – are conserved in all homologs from human herpesviruses (Supplementary Figure S3). The loop that sandwiches the outward-rotated nucleotides is less conserved; its first part (BALF2: Val923-Asn924; ICP8: Gly990-Met991) contacts the DNA in both structures, but additional interactions are only observed for ICP8, possibly thanks to additional stabilization provided by the other DNA strand in the dimer. Tyr937 is conserved as an aromatic residue in gamma- and betaherpesviruses, but Ser1005, its counterpart in HSV-1 ICP8, is likewise involved in DNA binding. Another secondary structure element with a conserved role in interacting with DNA is the β-strand that contains positively charged residues, which stabilize the phosphodiester backbone of the outward-rotated region.

Conversely, two regions interact with the DNA only in ICP8. The first of these is the helix spanning residues 787-801 which contacts nucleotides in positions −4 and −3. This element appears to be specific to DNA-binding proteins from alphaherpesviruses, as truncations in this region are observed in the other two families. The second region encompasses a mostly unstructured stretch of residues 543-576 which interact with the 3′ end of the visible DNA chain. Intriguingly, Tyr543 and Asp545, conserved throughout the herpesvirus family, do not interact with DNA in the BALF2-DNA structure due to the limited length of the modeled DNA strand.

In conclusion, our structural analyses showed conservation of key elements of DNA binding by ICP8 and BALF2, and sequence alignments suggest that these are universal features of herpesviral DNA-binding proteins.

### Structure of the ICP8 filament

ICP8 can form double helical protein-only filaments or coat ssDNA to produce single filaments^14,36^. To investigate the architecture of these filaments, we performed reconstitution experiments for both ICP8 and BALF2. We tested various DNA preparation methods and achieved optimal results using linear dsDNA with long ssDNA tails, similar to those used in the study of KSHV ORF6^37^. The linearized dsDNA was pretreated with T4 polymerase, which, in the absence of deoxynucleotide triphosphates, utilizes its 3′-5′ proofreading exonuclease activity, producing long single-stranded overhangs that can be coated by ICP8 or BALF2. This method yielded better defined filaments than those obtained using heat-denatured linear plasmid DNA (Fig. 4a). We attempted to generate protein-only filaments, but under identical conditions, incubation without DNA resulted in very short filaments (ICP8) or long filaments with complex trajectories (BALF2), limiting their suitability for cryo-EM studies.

**Figure 4.**
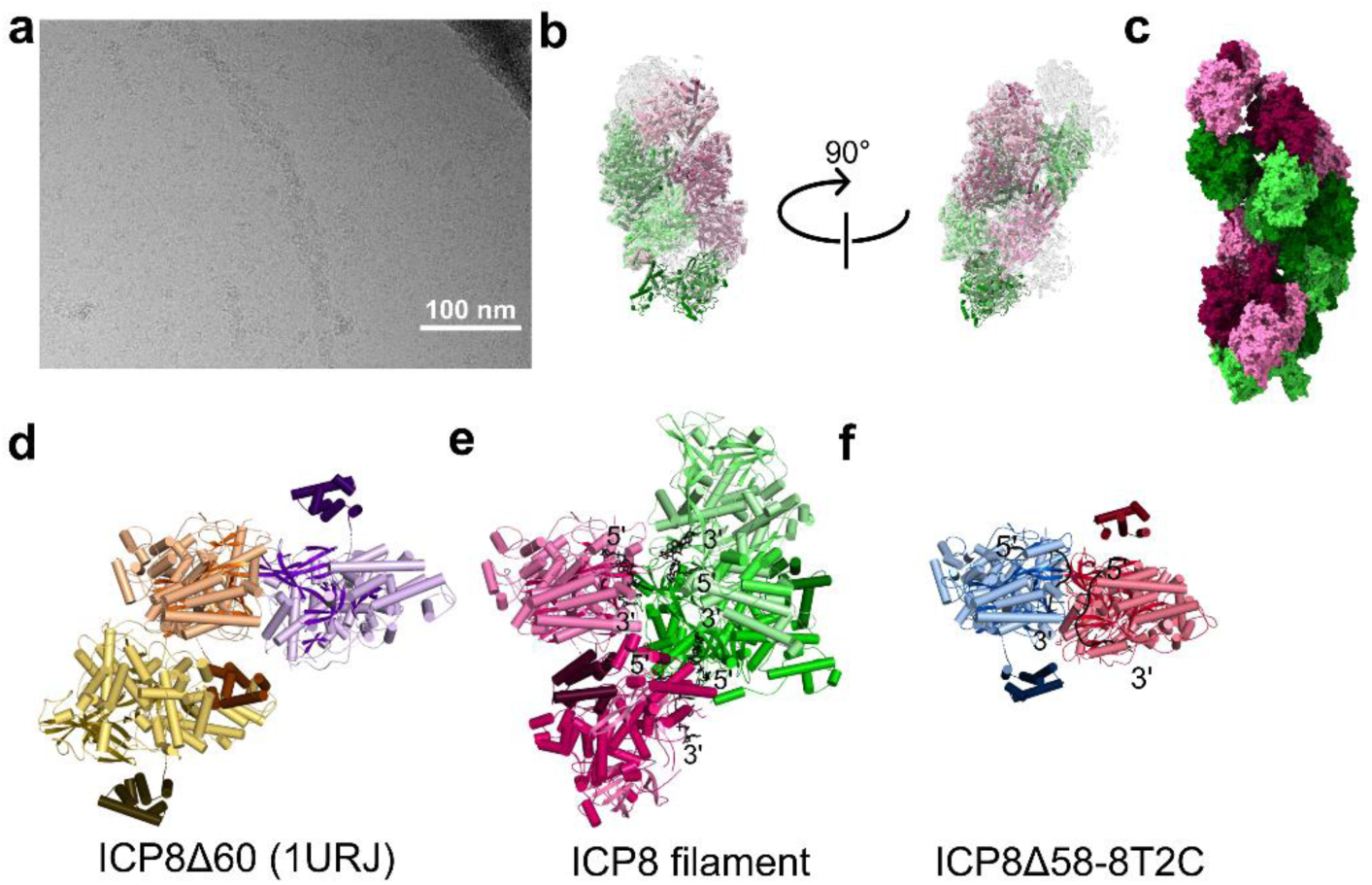
Cryo-EM structure of ICP8 filament and comparison with other ICP8 structures. **a)** Cryo-EM micrograph showing a double filament of ICP8. Scale bar represents 100 nm. **b)** Reconstruction of a segment of the double filament. The cryo-EM density map is shown in gray, and the modeled ICP8 subunits are shown in cartoon representation. **c)** Longer fragment of the filament modeled through superposition of multiple copies of the structure. ICP8 subunits are shown in surface representation and colored according to the strand they belong to (pink, green). **d-f)** Comparison of the arrangement of ICP8 subunits in different structures. Subunits are shown in cartoon representation, with C-terminal domains shown in darker shades. **d)** ICP8Δ58 apo crystal structure (PDB ID: 1URJ). Content of the asymmetric unit shown in orange and yellow and a single symmetry-related molecule in violet. **e)** Fragment of the ICP8 protein filament showing two interacting ICP8 subunits from different filament strands. Predicted positions of ssDNA fragments were modelled based on the crystal structure of the ICP8Δ58-DNA complex. **f)** ICP8Δ58-DNA crystal structure.

Cryo-EM data were collected for both BALF2-DNA and ICP8-DNA samples. 2D class averaging of BALF2-DNA data yielded results of insufficient quality for 3D reconstruction; however, observed classes indicated the presence of both single, protein-coated ssDNA strands and double helical filaments composed of antiparallel strands (Supplementary Figure S4).

Initial analysis of ICP8-DNA data using helical reconstruction proved unsuccessful due to the filaments not being linear. Instead, single-particle analysis was performed on selected filament regions. Ultimately, we calculated a reconstruction at 3.2 Å resolution, comprising a filament segment with five full-length protein chains and one chain representing only the C-terminal domain (Fig. 4b, Supplementary Figure S5). The filament exhibits a pitch of 267 Å and contains 6.62 subunits per turn, closely matching the helical parameters of previously characterized ICP8-only filaments^36^. The helical rise (Δ*z)* of the filament is 40.3 Å, and the helical twist is −54.4⁰. The filament is composed of two strands running antiparallel (Fig. 4c). Although DNA was used to produce the sample and appeared to enhance filament formation, it could not be built into the map. However, trace density was observed in areas corresponding to the DNA-binding site, as identified from our crystal structures. It is thus impossible to definitively determine whether the studied filament contains DNA, is composed solely of protein subunits, or the associated DNA has a partial occupancy.

An individual protein protomer within the filament makes contacts with four other full-length chains. Contacts with neighboring protomers within the same filament strand are formed in a head-to-tail arrangement, mediated through the C-terminal domains accommodated within a cleft of the N-terminal domain of the adjacent protein. This arrangement is identical to that observed in the crystal structures. However, within a single subunit, the C-terminal domain is positioned closer to the head domain compared to the arrangement in the crystal structure (Fig. 4d,e). Contacts are also formed with two protomers from the opposite antiparallel strand, with each interface exhibiting two-fold symmetry (Supplementary Figure S6). One interface is mediated by a loop (392-406), the end of a helix (691-694), and two residues from the following helix (713-714). Adjacent to this interface is the loop (989-996), which is involved in base-pairing stabilization in the crystal structure, and positioned near its counterpart from the other chain. The interface with the second protomer from the opposing strand involves the loop at the top of a hairpin (75-79, unmodelled in all crystal structures) and an unstructured fragment spanning residues 203-230.

While the reconstruction may correspond to a double protein-only filament, the binding of ssDNA to protomers can be modelled using the ICP8-DNA co-crystal structures (Fig. 4e,f). The modelled fragments within a single filament strand have the same polarity, suggesting a similar arrangement on long ssDNA is possible. However, the distance between modelled ssDNA fragments bound to protomers from the opposite filament strands is greater than that observed in the co-crystal structure, which likely represents the annealing step. This implies that if protein-DNA filaments of this morphology indeed form, they could represent an initial stage in the annealing process, requiring a rearrangement of the filament strands to bring the DNA strands closer and allow base-pairing.

### Interaction of ICP8 and UL12 at the ssDNA-dsDNA junction in the recombinase complex

To gain insights into the mechanistic properties of HSV-1 recombinase, we determined the cryo-EM structure of a reconstituted ternary complex of ICP8, UL12, and a model DNA at 3.29 Å resolution (Figure 5, Supplementary Figure S7). The DNA design included a double-stranded portion bound by UL12 and a 3′ ssDNA overhang to interact with ICP8, mimicking a result of UL12-mediated resection. The polypeptide chains are overall well-defined; however, the unstructured N-terminal portion of UL12 and the C-terminal domain of ICP8 are not resolved in the cryo-EM density. Map quality for the DNA was lower than for the protein part of the complex, and flexibility analysis of the reconstruction suggested increased mobility of the distal end of the dsDNA portion. Consequently, the DNA was built in two fragments – a dsDNA part with 9 base pairs and a 8-nt long ssDNA. The density for DNA between these two fragments is continuous, however not enough resolved to allow for DNA building.

**Figure 5.**
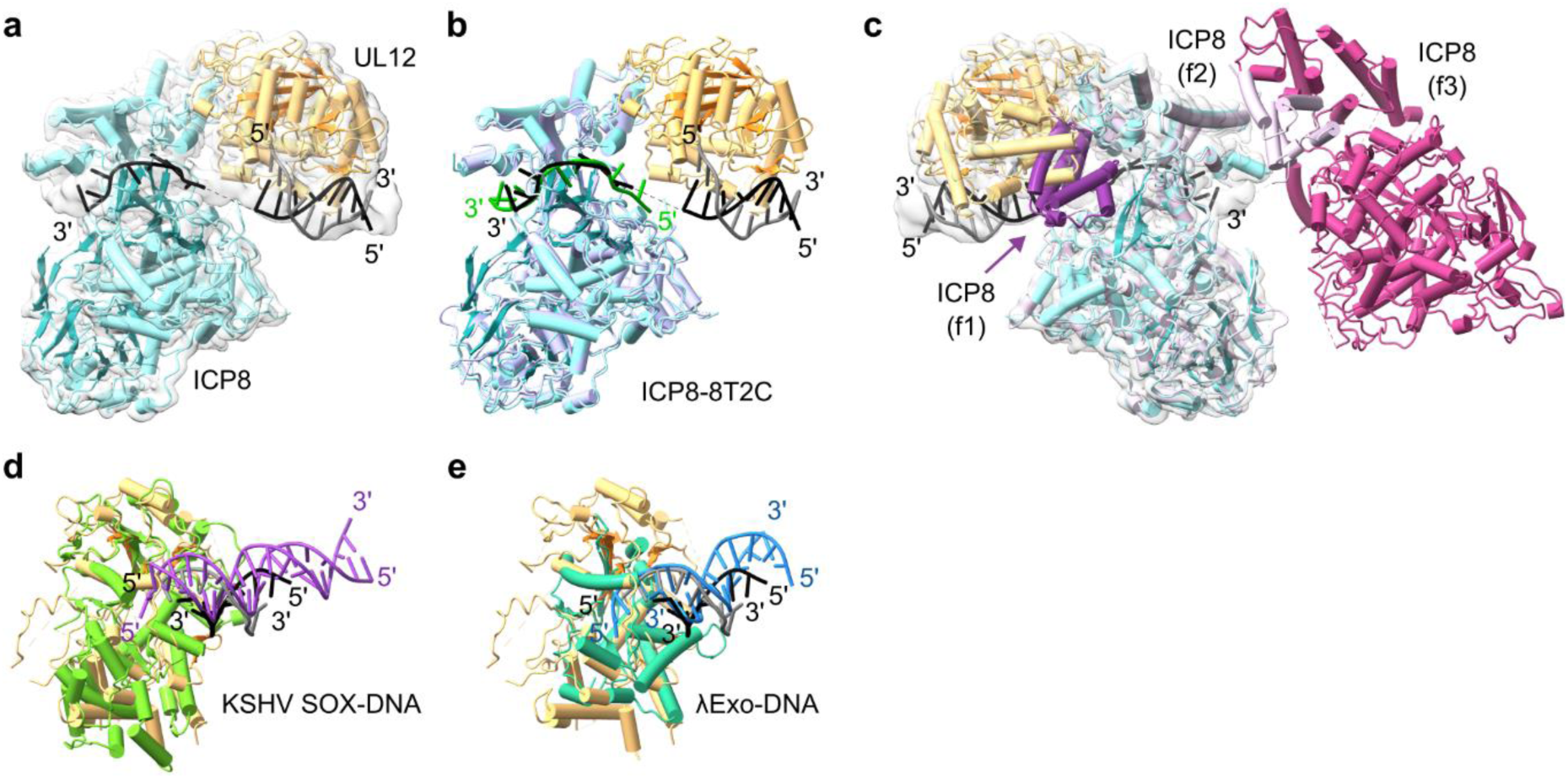
Structure of the HSV-1 recombinase in complex with DNA. **a)** Cryo-EM structure of the ICP8-UL12 complex bound to dsDNA with a long 3′ overhang. The cryo-EM density map is shown in gray. ICP8 and UL12 are shown in cyan and orange, respectively, with β-strands shown in darker shades. DNA strands are shown as black and gray ladder. **b)** Superposition of ICP8-DNA from the recombinase complex with a single ICP8-DNA complex from the co-crystal structure. The N-terminal domain of ICP8 from the crystal structure is shown in light blue and its bound ssDNA is shown in green. For clarity, the C-terminal domain is not displayed. **c)** Superposition of ICP8-DNA from the recombinase complex on the ICP8 subunits from the filament structure (three chains are shown, ICP8 f1-f3). The central subunit in this arrangement (light pink) was superimposed onto the ICP8 from the recombinase complex. The C-terminal and N-terminal domains from the adjacent subunits from the filament are shown in purple and pink, respectively. **d)** Superposition of dsDNA complexes of HSV-1 UL12 and KSHV SOX (PDB ID: 3POV). KSHV SOX is shown in bright green, and the dsDNA bound to it is shown in violet. **e)** Superposition of dsDNA complexes of HSV-1 UL12 and λExo (PDB ID: 3SM4). λExo is shown in teal, and the dsDNA bound to it is shown in blue.

In the complex, ICP8 binds to the ssDNA portion as it emerges from UL12 (Fig. 5a). The interaction between ICP8 and UL12 buries a surface area of 939.2 Å^2^. One side of the head domain of ICP8 interacts mainly with the N-terminal lobe of UL12. The DNA strand is transferred from UL12 to ICP8 without significant trajectory changes. The ssDNA conformation and its interaction with ICP8 are similar to those observed in the co-crystal structures (Fig. 5b). The ICP8 surface required for the formation of the dimer observed in our crystal structures (potentially corresponding to the DNA annealing step) remains available. Conversely, the position occupied by UL12 partially overlaps with the interaction site for the C-terminal domain of another protomer in a filament (Fig. 5c). However, on a longer 3′ overhang, the ICP8 in this complex could use its C-terminal domain to interact with another ICP8 protomer bound to the longer ssDNA (Fig. 5c). This implies that ssDNA generated by the nuclease can be directly coated by ICP8 forming protein-DNA filament.

The binding mode of UL12 to dsDNA is similar to that of its KSHV homolog, SOX (Fig. 4d). The 5′ recessed end of the DNA is positioned in the vicinity of the active site, but it does not reach it. Its slightly different position compared to the SOX-dsDNA structure can be attributed to the absence of divalent ions in the sample, intended to prevent DNA degradation. Furthermore, for a productive interaction with the active site, a 5′ phosphate group would be required, as illustrated by the structures of λExo^38^ (Fig. 5e). The arrangement observed in our complex closely resembles the nonproductive complex of λExo with a blunt-ended dsDNA with a 5′-hydroxyl group. This is why we decided to assign the nucleotides in the visible part of the duplex to positions 1–9 of the 12-nt strand. In our structure, the bridge loop (residues 259– 271), which spans across the cleft that contains the active site, is fully modeled. The helix that follows it, abuts the last base pair, and the sidechain of Val273 contacts the base and the ribose of the 5′-terminal nucleotide of the cleaved strand. The sidechain of Lys407 interacts with the non-bridging oxygen in the phosphate group between the second and third nucleotide of the cleaved strand. Contacts are also made on the other side of the major groove with the complementary strand of the duplex. These contacts are mediated by the sidechain and backbone of Arg443, and the sidechain of Arg496.

In conclusion, we provide the first structural information for a nuclease-DNA-binding protein complex involved in recombination which shows the handoff of the DNA between the two proteins.

## DISCUSSION

The DNA-binding proteins are essential players in herpesvirus DNA replication. ICP8 has been shown to be involved in each step of the process: i) establishment of prereplicative sites, ii) assisting in replication initiation, iii) recruiting other components or stimulating their activity within nuclear replication compartments, and finally iv) promoting genomic recombination, which leads to the formation of branched DNA structures^1^. In our study, we provide structural insights into how HSV-1 ICP8 and EBV BALF2 interact with DNA, shedding light on the mechanistic details of their functions.

We determined high-resolution structure of BALF2 in DNA-bound form. Within the complex, the nucleic acid is bound within a positively charged cleft between the head and shoulder subdomains. The DNA interacts with the protein mainly via the bases; however, a segment of four nucleotides has an altered orientation, with the base edges exposed to the solvent. We also analyzed the affinity of different BALF2 variants for a 14-mer DNA oligonucleotide to assess the role of individual residues. Truncation of the C-terminal fragment of BALF resulted in a fourfold decrease in DNA binding. This contrasts with findings for the BALF2 homolog from KSHV, termed ORF6, which exhibited similar affinity with both the full-length and a C-terminally truncated variant (K_d_ of 220 µM and 150 µM, respectively)^37^. How the presence of this extreme C-terminal fragment affects binding to a short ssDNA fragment, with the length of a footprint of single protein subunit, remains unclear.

We also present two crystal structures of ICP8 complexes with 14-mer DNA oligonucleotides. Regions involved in DNA binding, that were unmodelled in the previously published apo structure, can now be visualized: the 989-997 loop that stabilizes the outward-rotated bases, the 787-801 helix that interacts the DNA upstream, and the partially helical region spanning residues 554-572 that contacts the 3’ distal part of the DNA. The two-fold symmetric dimer found in the asymmetric unit of the ICP8-DNA crystal structure presented herein has a similar arrangement to the one observed between in the apo ICP8 structure between symmetry-related molecules^28^. However, in the presence of DNA the complex becomes tighter. One of the contacts uniquely present in the complex with DNA is formed between the sidechains of Val760 and Gln706. Remarkably, Q706A substitution resulted in a complete loss of strand annealing while not affecting other functions of ICP8^39^. This implies that the conformation of the two-fold symmetric ICP8 dimer observed in our structures is essential for DNA annealing.

A striking feature of the DNA strands captured in the complex with ICP8 is the trajectory of their backbones. Most nucleobases face the protein surface, with the exception of a stretch of three or four nucleotides that are rotated outward, positioning them away from the protein and towards the similarly exposed nucleobases of the DNA bound by the other ICP8 protomer. Nucleobases from the two strands are thus presented to each other, enabling base-pairing between them. Depending on the DNA oligonucleotide used for co-crystallization, we observed different patterns of base-pair formation – paring between non-complementary bases in all positions, or formation of two Watson-Crick base pairs but with a one-nucleotide register shift. The non-Watson-Crick pairings likely result from the sequences of the DNAs used. Analysis of all our structures indicates that ICP8 has a preference for exposing bases located downstream of a purine nucleotide. This, together with the stabilizing effect of at least partial complementarity between the exposed regions, may determine the register with which ICP8 interacts with the oligonucleotide. Overall, our structures show a high degree of flexibility in base-pair formation, which could be important during homology search.

The outward rotation of nucleotide bases has already been proposed as an element of the ICP8 strand invasion mechanism^20^. A very similar change in backbone trajectory is apparent in our cryo-EM structure of BALF2. To date, the proposed roles of BALF2 have included melting of secondary structures on the DNA replicated by the polymerase, which reduces Pol pausing and increases product length,^33^ and destabilization of the DNA helix^34^. The ability of BALF2 to form two-fold symmetric dimers could allow it to play a role in DNA annealing, similar to ICP8.

Our next aim was to understand the architecture of filaments formed by ICP8 and BALF2. The propensity to form double filaments was previously shown for ICP8, but also for KSHV ORF6^35,37^. In our experiments, we used a similar type of DNA as in the ORF6 study and observed that the addition of DNA increased the efficiency of filament formation. However, when we prepared a reconstruction based on the cryo-EM data for the ICP8 filament, we noticed that its helical parameters were very similar to those estimated for the protein-only ICP8 filaments reported earlier^36^. However, ill-defined density located at the DNA-binding site of ICP8 was observed, suggesting the DNA could still be present there. This is further supported by the fact that the double helical protein-only filaments of ORF6 were shown to be able to envelop ssDNA tails without altering the filament architecture^37^.

Our high-resolution reconstruction of the ICP8 filament allowed us to better define the interprotein interactions required for filament formation. We were able to describe the interfaces between subunits located on the opposing strand of the double filament. An interaction between two conserved motifs – FNF (Phe1142-Asn1243-Phe1144), located in the extreme C-terminus of the protein, and FW (Phe843-Trp844) – is essential for the formation of the double filament of ICP8^4^. Even though the full-length ICP8 form was used for filament reconstitution, the extreme C-terminus of the protein could not be accurately modelled. However, the last residues that could be resolved (1134/1135) come very close to the tip of the head domain of the neighboring protomer, approaching the helix containing the FW motif from the other side. Remarkably, a patch of unoccupied map density is consistently observed in all modelled subunits in the area where the FNF-FW interaction is expected to form, in the vicinity of residues 850, 854, 862, 869. It should be noted that there is no overlap between the regions of ICP8 that engage in protein-protein contacts in the double filament and the regions involved in the interaction of ICP8 with ssDNA or two-fold symmetric protein-protein interface formed during DNA annealing. This implies that the formation of a double filament containing two antiparallel DNA strands could be possible, which could then undergo a rearrangement whereby the two DNA strands would be brought together, resulting in the supercoiling of the filament.

For the BALF2-DNA filament sample, two types of morphologies could be observed. One appeared to correspond to single filaments with a similar appearance to the assemblies observed in the presence of 90 nt DNA. The other 2D classes were very similar to the double filaments observed in the ICP8-DNA sample. This indicates that BALF2 is capable of forming double helical filaments, but further analyses will be required to confirm their molecular architecture.

In dsDNA viruses whose DNA replication mechanisms involve recombination events, single-strand annealing proteins (also termed annealases) are found to cooperate with nucleases, forming so-called Exonuclease-Annealase Two-component Recombinase systems (EATRs)^27^. Well-studied examples of such systems are the λExo nuclease and Red β annealase of phage lambda and RecE exonuclease and RecT annealase of Rac prophage. In this study, we present the first complete structure of a recombinase complex containing both proteins and a model DNA. Similarly to λExo and RecE, UL12 is a PD-(D/E)XK family nuclease; however, unlike them, it functions as a monomer. Another intriguing similarity regards the site of exonuclease interaction with the annealase. The C-terminal domain of Red β interacts with λExo using a site that partially overlaps with that to which the bacterial single-stranded DNA-binding protein (SSB) binds. This mutually exclusive binding was proposed to be the molecular basis of a ‘hand-off’ mechanism, in which the Red β, loaded onto the single strand of DNA exposed by the exonuclease action of λExo, can next direct it to an SSB-coated DNA fragment to initiate complementarity search. In HSV-1, ICP8 performs the role of both the DNA-binding protein and the annealase. UL12 binds near its concave surface, very close to the site where the C-terminal domain of an adjacent ICP8 molecule would bind. It is thus possible that competition for this binding site between the nuclease and another ICP8 molecule is also used as a mechanism of directing the exposed ssDNA to initiate recombination e.g. at the site of a collapsed replication fork.

An important application of EATRs is their use in recombination-mediated genetic engineering, or recombineering^40^. In this approach, homologous recombination executed by EATR systems is leveraged to enable efficient and precise editing of the DNA sequence *in vivo*, even in large genetic constructs. Both the λExo-Red β and RecET system were used for this purpose; however their efficacy was largely limited to *E. coli* and cognate species^41^. A likely reason for this is the fact that the EATR complexes cooperate with cellular proteins, making these systems host-specific. The HSV-1 recombinase complex has already been postulated as a promising tool with the potential to introduce recombineering into human cells^42^. The structure of the HSV-1 recombinase complex offers valuable insight into the mechanistic details of the initial step of recombination paving the way to engineering this system for improved performance in genetic engineering.

Various functional and structural features of ICP8 are shared with different types of proteins involved in ssDNA transactions. It is an ssDNA-binding protein that holds the DNA strand in an extended conformation, but it has also been classified as an annealase thanks to its capability of performing strand transfer between homologous strands of DNA. Similarly to classic recombinases, it is found to form helical filaments both with and without DNA^43^. The structures presented herein allow us to identify intriguing similarities and differences in the way ICP8 and BALF2, and functionally related proteins interact with DNA. A common feature among ssDNA-interacting proteins is the formation of homooligomeric rings. Such rings or gapped ellipses have been observed for Rad52, Redβ, DdrB, but also for ICP8^43,44^. Another recurring feature is the organization of the bound ssDNA into shorter segments. The length of these segments varies for different proteins and is usually correlated to the size of their footprints for DNA binding. In the Rad51/RecA recombinase family, these are nucleotide triplets sandwiched between two flanking loops^45^. This is similar to the stabilization observed in our ICP8-DNA structures, where the 989-997 loops from each of the ICP8 protomers sandwich paired triplets. In human RAD52, the ssDNA strand is divided into stretches of four nucleotides^46^. In the structure of a filament of Red β homolog from a Listeria prophage with a dsDNA annealing intermediate, a β-hairpin wedges into the duplex every 5 base pairs^47^. The structure of a protein-DNA filament of Ha-LEF3, an SSB from *Helicovera armigera* nucleopolyhedrovirus, shows a disruption of base stacking every 6 nucleotides, in the middle of the binding site of a single protomer^48^.

In all the abovementioned cases, the bound ssDNA is stabilized mostly through interactions with the phosphate backbone, which allows the nucleobases to be presented outwards, thus enabling homology search and base-pairing with a complementary strand. However, even though both ICP8 and BALF2 have a footprint of ∼15 nucleotides, in their complex structures, the interaction with the DNA is mediated mostly by the bases, with only a short stretch of the ssDNA coordinated via the phosphate backbone. Intriguingly, a similar situation is observed in the DNA-complex of DdrB, which is a *D. radiodurans* single-strand annealing protein^49^. In this complex, each subunit of the homopentamer interacts with 6 nucleotides, but only in four of them are the bases exposed outwards, and the other two bases remain buried through interaction with the protein. This allows for an elegant two-step mechanism of ssDNA annealing. At first, two DNA-bound pentamers associate in a transient manner, allowing for base-pairing only between a subset of DNA bases. If complementarity is found, the increased torsional strain imposed on the sugar phosphate backbone caused by the base-pairing destabilizes the protein-DNA interaction, releasing the buried bases and enabling full base-pairing. Remarkably, the protein-protein contacts in the ICP8-DNA crystal structure are also limited, suggesting that DNA conformation-driven release of ICP8 upon base-pairing could also be happening here.

DNA-binding proteins are essential elements of the herpesviral DNA replication machinery and our studies provide structural information about the architecture of several important complexes formed by these proteins during various steps of DNA replication and recombination, including ssDNA coating, DNA strand annealing, and annealase loading onto ssDNA exposed by nuclease action. We are also presenting a high-resolution structure of the double filament of ICP8, revealing the protein-protein interactions that enable its assembly. Our structural work provides a framework for future studies and the complete understanding of the dynamic steps of viral recombination.

## Supporting information

Suuplementary Figures S1-S7

## Acknowledgements

We would like to thank Iwona Ptasiewicz for technical assistance, and S. Diane Hayward and Mikhail Alexeyev for sharing the genetic constructs. We acknowledge DESY (Hamburg, Germany), a member of the Helmholtz Association HGF, for the provision of experimental facilities. Parts of this research were carried out at PETRA III and we would like to thank Guillaume Pompidor for assistance in using P11 beamline. Beamtime was allocated for proposal BAG-20230001 EC. This work was financed by the OPUS grant from the National Science Center, Poland (2022/45/B/NZ1/02456). This publication was developed under the provision of the Polish Ministry of Education and Science project, ‘Support for research and development with the use of research infrastructure of the National Synchrotron Radiation Centre SOLARIS,’ under contract no. 1/SOL/2021/2. We acknowledge the SOLARIS Centre for access to the cryo-EM Beamline, where the measurements were performed. DNA constructs for the generation of recombinant baculoviruses insect cell cultures for recombinant protein production were generated, respectively, at the IIMCB Genomic Engineering and Preclinical Drug Development core facilities (IN-MOL-CELL Infrastructure) funded by the European Union – NextGenerationEU under National Recovery and Resilience Plan. IN-MOL-CELL Infrastructure was also funded by the European Union under Horizon Europe (Project 101059801 - RACE) and by RACE-PRIME project carried out within the IRAP programme of the Foundation for Polish Science co-financed by the European Union under the European Funds for Smart Economy 2021-2027 (FENG).

## Author contributions

G.R.A. purified ICP8 and BALF2 proteins, performed the structural studies and determined the structures of BALF2 and ICP8 with DNA. S.P. purified ICP8 and UL12 proteins, reconstituted the ICP8-UL12-DNA complex and determined its structure. G.R.A., S.P. and M.C.C. analyzed cryo-EM data. A.J. purified BALF2 variants. J.J. produced recombinant proteins in insect cell cultures. J.P. analyzed BALF2 DNA-binding data. M.N. and M.F. supervised the project and wrote the manuscript with input from all coauthors.

## Notes

### Competing Interest Statement

The authors have declared no competing interest.

