## Supplementary material for "Structural insights into DNA annealing and recombination by herpesviral DNA-binding proteins ICP8 and BALF2": Suuplementary Figures S1-S7

### SUPPLEMENTARY INFORMATION

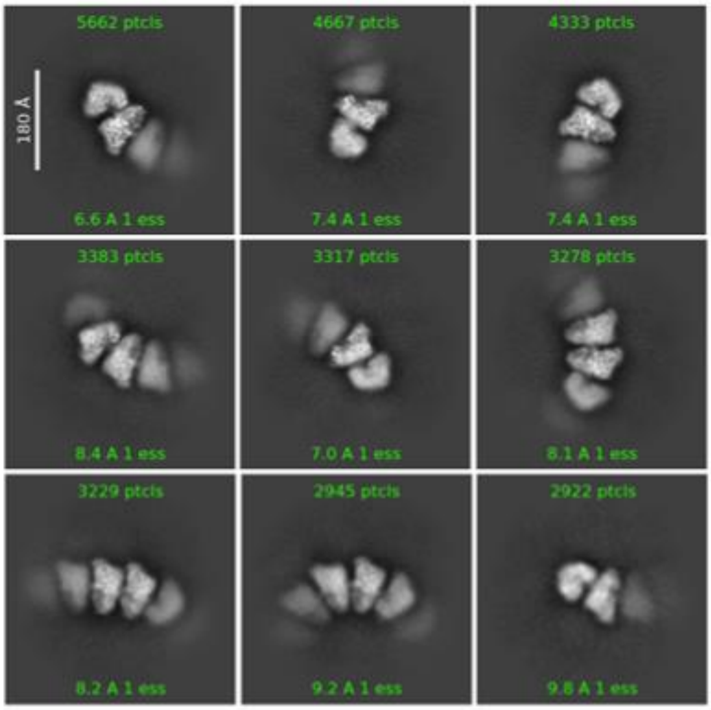


Supplementary Figure S1. **Representative 2D classes observed during the processing of the BALF2-90mer data.** The classes are the averages of particles extracted with a size of 256 pix which corresponded to 430 Å.

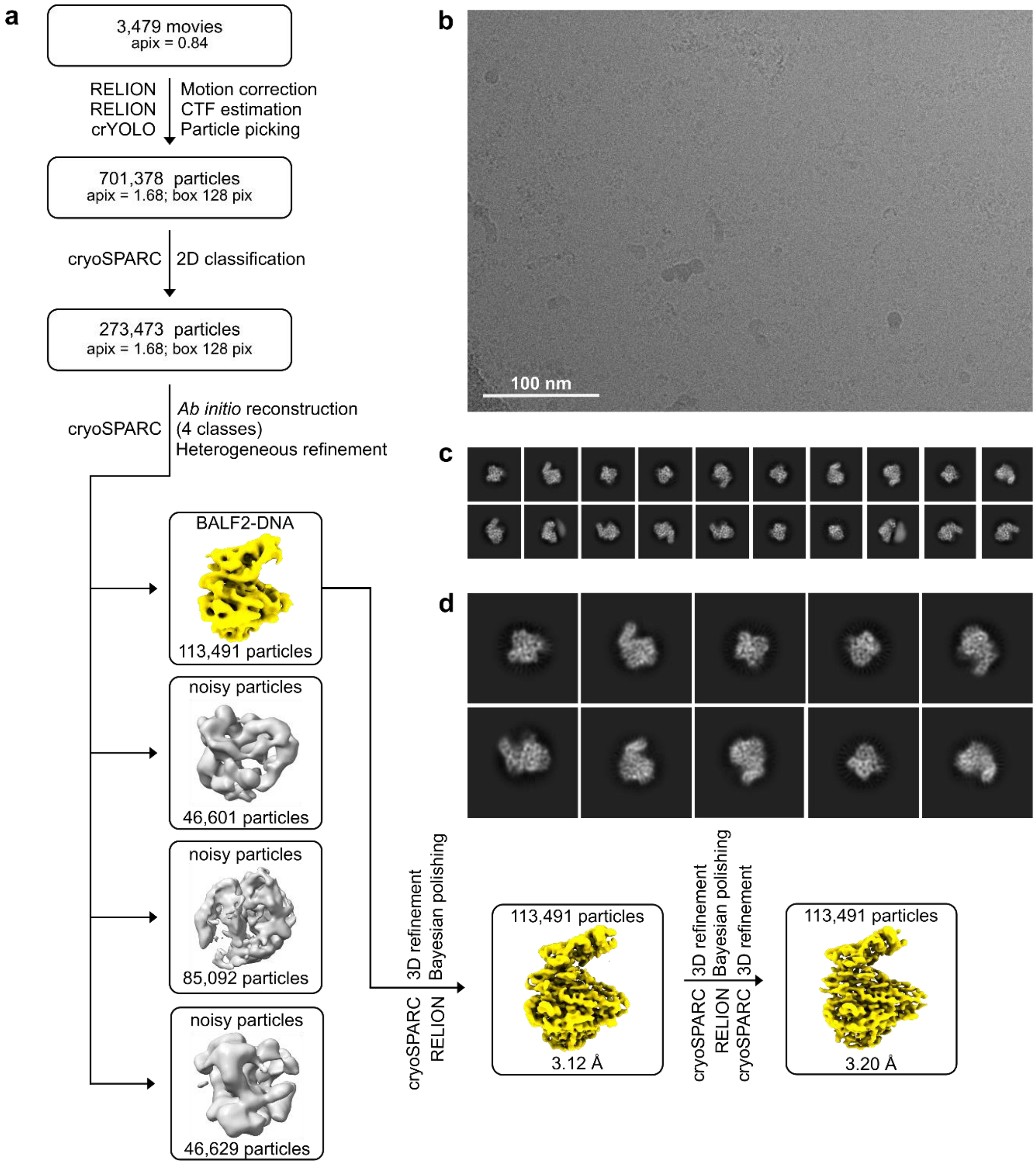


Supplementary Figure S2. **BALF2-ssDNA cryo-EM data processing. a)** Three-dimensional reconstruction pipeline. **b)** Representative micrograph. **c)** Class averages from the 2nd round of 2D classification in cryoSPARC. **d)** Enlarged view of class averages from the 2nd round of 2D classification in cryoSPARC.

```
HSV1        1  -METKPKTATT---IKVPPGPLGYVYARACPSEGIEL--LALLSARSGDSDVAVAPLVVGLTVESGFEANVAVVVGSRTTGLGGT  79
HSV2        1  -MDTKPKTTTT---VKVPPGPMGYVYGRACPAEGLEL--LSLLSARSGDADVAVAPLIVGLTVESGFEANVAAVVGSGTTGLGGT  79
VZV         1  -MENTQKTVT------VPTGPLGYVY--ACRVEDLDLEEISFLAARSTDSDLALLPLMRNLTVEKTFTSSLAVVSGARTTGLAGA  76
HHV-6A      1  ---MADENETV---VSAPVSTAAWIYVFPKDKELLDV--LSVLSLMERNSPIVISPLLMNLTVENDFSTTVKTPITNF-------  70
HHV-7       1  ---MADDNETV---VSAPICTAAWLYILPKEQKLIEI--LTTLSLMEKRKSVVISPLLLNLTVENDFFPTVKTPIINY-------  70
HCMV        1  ----MSHEELT---ALAPVGPAAFLYFSRLNAETQEI--LATLSLCDRSSSVVIAPLLAGLTVEADFGVSVRTPVLCY-------  69
KSHV        1  MALKGPQTLEENIGSAAPTGPCGYLYAYLTHNFPIGE--ASLLGNGYPEAKVFSLPLLHGLTVESDFPLNVKAVHKKID------  77
EBV         1  --MQGAQTSEDNLGSQSQPGPCGYIYFYPLATYPLRE--VATLGTGYAGHRCLTVPLLCGITVEPGFSINVKALHRRPD------  75
Cons_ss:                           eeeeeeee    hhh hhhhh          eeeeee   ee      eeeeeee

HSV1       80  AVSLKLTPSHYSSSVYVFHGGRHLD-PSTQAPNLTRLCERARRHFGFSDYTPRPGDLKHETTGEALCERLGLDPDRALLYLVVTE  163
HSV2       80  AVSLKLMPSHYSPSVYVFHGGRHLA-PSTQAPNLTRLCERARRHFGFSDYAPRPCDLKHETTGDALCERLGLDPDRALLYLVITE  163
VZV        77  GITLKLTTSHFYPSVFVFHGGKHVL-PSSAAPNLTRACNAARERFGFSRCQGPPVDGAVETTGAEICTRLGLEPENTILYLVVTA  160
HHV-6A     71  GGTILTKITSFMPVCFFFHGTEQLVGMAEDHGDLIRLCEQTRQKFHLQSFEVPTAR--KVIDIKALCSAVGKDADSVICHVACGN  153
HHV-7      71  GGTVITKITSFMPVCFFFHGTDVFLKEAEDHGNLDKLCKQTREKFNLQEFVVNGNR--KSVDIGKICESVGRNADDVLCHIVVGN  153
HCMV       70  DGGVLTKVTSFCPFALYFHHTQGIVAFTEDHGDVHRLCEDARQKYALEAYMPEADR--VPTDLAALCAAVGCQASETTVHVVVGN  152
KSHV       78  ATTASVKLTSYHREAIVFHNTHLFQ-PIFQGKGLEKLCRESRELFGFSTFVEQQ-HKG-TLWSPEAC-PQLPCANEIFMAVIVTE  158
EBV        76  PNCGLLRATSYHRDIYVFHNAHMVP-PIFEGPGLEALCGETREVFGYDAYSALP-RE--SSEPGDFF-PEGLDPSAYLGAVAITE  155
Cons_ss:         eeeeeeee  eeeeee                hhhhhhhhhhhh                hhhhhhh    hhhhhhhhhhhh

HSV1      164  GFKEAVCINNTFLHLGGSDKVTIGGAEVHRIPVYPLQLFMP-----DFSRVIAEPFNANHRSIGEKFTYPLPFFNRPLNRLLFEA  243
HSV2      164  GFREAVCISNTFLHLGGMDKVTIGDAEVHRIPVYPLQMFMP-----DFSRVIADPFNCNHRSIGENFNYPLPFFNRPLARLLFEA  243
VZV       161  LFKEAVFMCNVFLHYGGLDIVHINHGDVIRIPLFPVQLFMP-----DVNRLVPDPFNTHHRSIGEGFVYPTPFYNTGLCHLIHDC  240
HHV-6A    154  GFKELLFAGLLIPCVEEQIQVQVGEYSCVKIPLYSATLFET------EETISLSSCTEF--------IQERGFFLPALSETLFYY  224
HHV-7     154  GFKELLFAGLLIPCVEEQIQVQVGECLAIKIPLYSATLFES------EETLCIDTCTEF--------IQENGFYAPQISEVLFYL  224
HCMV      153  GLKEFLFAGQLIPCVEEATTVRLHGGEAVRVPLYPPTLFNSLQLDAEADEVSLDARSAF--------VEARGLYVPAVSETLFYY  229
KSHV      159  GFKERLYGGKLVPVPSQTTPVHIGEHQAFKIPLYDEDLFGP----------------------SRAQELCRFYNPDISRYLHDS  220
EBV       156  AFKERLYSGNLVAIPSLKQEVAVGQSASVRVPLYDKEVFPE----------------------GVPQLRQFYNSDLSRCMHEA  216
Cons_ss:       hhhhhhhh  eeee    eeeee  eeeeee    hhh                                      hhhhhhhhhh

HSV1      244  VVGPAAVALRCRNVDAVARAAAHLAFDENHEGAALPADITFTAFEASQG--KTPRGGR-----DGGGKGAAGGFEQRLASVMAGD  321
HSV2      244  VVGPAAVALRARNVDAVARAAAHLAFDENHEGAALPADITFTAFEASQG--KPQRGAR-----DAGNKGPAGGFEQRLASVMAGD  321
VZV       241  VIAPMAVALRVRNVTAVARGAAHLAFDENHEGAVLPPDITYTYFQSSSSGTTTARGARRNDVNSTSKPSPSGGFERRLASIMAAD  325
HHV-6A    225  VFTSWGTTLRFSNTKELIDAGLKQFTQDGEQTVKLAPHKTYLGISGQKI-------------------SAVEKDFLMLVDSVVTE  290
HHV-7     225  IFTSWGMTLRFNNTLELIKAGLKQFIQDTEQTVKLAPNKTYHGIPGQKL-------------------SPIEKDHLMLVDAVITE  290
HCMV      230  VYTSWCQSLRFSEPRVLIEAALRQFVHDSQQSVKLAPHKRYLGYMSQRL-------------------SSLEKDHLMLSDAVVCE  295
KSHV      221  IFTGIAQALRVKDVSTVIQASERQFVHDQYKIPKLVQAKDFPQCAS---------------------RGTDGSTLMVIDSLVAE  283
EBV       217  LYTGLAQALRVRRVGKLVELLEKQSLQDQAKVAKVAPLKEFPASTI---------------------SHPDSGALMIVDSAACE  279
Cons_ss:       hhhhhhheeee hhhhhhhhhhhhhh  eeee                                        hhhhhhhhhhh

HSV1      322  AALALESIVSMAVFDEP-PTDISAWPLFEGQDTAAARANAVGAYLARAAGLVGAMVFSTNSALHLTEVDDAGP----ADPKDHSK  401
HSV2      322  AALALESIVSMAVFDEP-PPDITTWPLLEGQETPAARAGAVGAYLARAAGLVGAMVFSTNSALHLTEVDDAGP----ADPKDHSK  401
VZV       326  TALHAEVIFNTGIYEET-PTDIKEWPMFIGTEGTLPRLNALGSYTARVAGVIGAMVFSPNSALYLTEVEDSGM----TEAKDGGP  405
HHV-6A    291  LSFSHVAEYLDSVYDPSQIMNFNDWPIIRNSETHAERMAQLTNLKLHLSSHLAVLIFAPNSILYCSKLAFIPNVK--QAFN-SVM  372
HHV-7     291  LTFSYTAEYLDSIYENNQIMNFSEWPIIKSAETHEEKIVELKKLRLHLSSHVAALVFAANSILYSNKLAYISNTK--QAFN-SAI  372
HCMV      296  LAFSFASVFFDSAYQPAESMLFSEWPLVTNATDHRDLIRALTELKLHLSTHVAALVFSANSVLYQHRLVYLQSSARHPSAG-GTA  379
KSHV      284  LGMSYGLSFIEGPQDSCEVLNYDTWPIFENCETPDARLRALEVWHAEQALHIGAQLFAANSVLYLTRVAKLPQ----KNQRGDA-  363
EBV       280  LAVSYAPAMLEASHETPASLNYDSWPLFADCEGPEARVAALHRYNASLAPHVSTQIFATNSVLYVSGVSKSTG-------QGKE-  356
Cons_ss:       hhhhhhhhhh                      hhhhhhhhhhhhhhhhhhhhhhhh   eeeeee

HSV1      402  ---PSFYRFFLVPGTHVAANPQVDREGHVVPGFEGRPTAPLVGGTQEFAGEHLAMLCGFSPALLAKMLFYLERCDGAVIVGRQEM  483
HSV2      402  ---PSFYRFFLVPGTHVAANPQLDREGHVVPGYEGRPTAPLVGGTQEFAGEHLAMLCGFSPALLAKMLFYLERCDGGVIVGRQEM  483
VZV       406  G--PSFNRFYQFAGPHLAANPQTDRDGHVL---SSQSTG---SSNTEFSVDYLALICGFGAPLLARLLFYLERCDAGAFTGGHG-  481
HHV-6A    373  TQELLLRSLSFCNALSSLSDDVYNDNRKIIKCDSTS------GKDDKFSANHLAYACATSPQLLSFVVWNLNRMSVYNAGN-AHT  450
HHV-7     373  TQETLLRSIQFCNSLSSLNEDFYNDARKLIKCNSSP------CKEDKFSAFHLAYACATCPQILSHIIWNLNRMSIYNTNC-GNS  450
HCMV      380  SQETLLKAIQFTNGLSAACEDVYNDARKVLKFQGAP------LKDERYGPQHLALVCGTCPQLVSGFVWYLNRVSVYNTGLSGSS  458
KSHV      364  ---NMYNSFYLQHGLGYLSEATVKENGASAFKGVPVSAL----DGSSYTLQHLAYASSFSPHLLARMCYYLQFLPHHKNTNSQSY  441
EBV       357  ---SLFNSFYMTHGLGTLQEGTWDPCRRPCFSGWGGPDVTGTNGPGNYAVEHLVYAASFSPNLLARYAYYLQFCQGQKSSLTPVP  438
Cons_ss:             eeeehh  hh    hhhhh                           hhhhhhhhh hhhhhhhhhhhhhh

HSV1      484  DVFRYVADSNQTDVPCNLCTFDTRHACVHTTLMRLRARHPKF-ASAARGAIGVFGTMNSMYSDCDVLGNYAAFSALKRAD-----  562
HSV2      484  DVFRYVADSGQTDVPCNLCTFETRHACAHTTLMRLRARHPKF-ASAARGAIGVFGTMNSAYSDCDVLGNYAAFSALKRAD-----  562
VZV       482  DALKYVTGTFDSEIPCSLCEKHTRPVCAHTTVHRLRQRMPRF-GQATRQPIGVFGTMNSQYSDCDPLGNYAPYLILRKPGD----  561
HHV-6A    451  EIYNHLVNCSANL--CEFCDGKCCQSCIGTAMVRVGTRLPAIPKNVKKEPLVMSMF-SRYYAEVDILGSFGRKPVSELKEIG---  529
HHV-7     451  EIYNHIVNCSSNL--CEFCEGKCCHSCIGTALIRINSRLPQISKTTKKEPIVMTMF-SRFYADVDVLGSFGKKGVNESKDPM---  529
HCMV      459  TLTNHLVGCAAGL--CEACGGTCCHTCYQTAFVRVRTRLPVVPKQPKKEPCVITVQ-SRFLNDVDILGSFGRRYNVDAKDGGLDG  540
KSHV      442  NVVDYVGTAAPS-QMCDLCQGQCPAVCINTLFYRMKDRFPPVLSNVKRDPYVITGTA-GTYNDLEILGNFATFREREEEG-----  519
EBV       439  ETGSYVAGAAAS-PMCSLCEGRAPAVCLNTLFFRLRDRFPPVMSTQRRDPYVISGAS-GSYNETDFLGNFLNFIDKEDDG-----  516
Cons_ss:        hhhhhhhhh     hhh     hhhhhhhhhhhhhh                eeeee

HSV1      563  ---------------------------------------------GSETARTIMQETYRAATERVMAELETLQYVDQAVPTAMGR  602
HSV2      563  ---------------------------------------------GSENTRTIMQETYRAATERVMAELEALQYVDQAVPTALGR  602
VZV       562  ---------------------------------------------QTEAAKATMQDTYRATLERLFIDLEQERLLDRGAPCSSEG  601
HHV-6A    530  ---------------------------------------------KDQQN-TLS-LDRGKFVSQIFDYCKKNSLIDPVTG--EDT  565
HHV-7     530  ---------------------------------------------KEAQT-TPS-LDRFKFLGMIHDYCKKNNLIDAITG--EDN  565
HCMV      541  KGDDGVPGGGAGGGGGRDVSGGPSDGLGGGRGGGGGGDSGGMMGRGGRML-GAS-VDRTYRLNRILDYCRKMRLIDPVTG--EDT  621
KSHV      520  ---------------------------------------------NPVED-APK-YTYWQLCQNITEKLASMGI--SEGG---DA  552
EBV       517  ---------------------------------------------QRPDD-EPR-YTYWQLNQNLLERLSRLGI-DA-EG---KL  549
Cons_ss:                                                           hhhhhhhhhhhhhhh
```

```
HSV1       603  LETIITNREALHTVVNNVRQVVDREVEQLMRNLVEGRNFKFRDGLGEANHAMSLTLDPYACGPCPLLQLLGRRSNLAVYQDLALS  687
HSV2       603  LETIIGNREALHTVVNNIKQLVDREVEQLMRNLIEGRNFKFRDGLAEANHAMSLSLDPYTCGPCPLLQLLARRSNLAVYQDLALS  687
VZV        602  LSSVIVDHPTFRRILDTLRARIEQTTTQFMKVLVETRDYKIREGLSEATHSMALTFDPYSGAFCPITNFLVKRTHLAVVQDLALS  686
HHV-6A     566  F--NVRSKKDFVSIIHGLTQCIEECVSRCIVEMRR-TQT-PREQIENCLQSFNVDTTPYATAFSPFLTFSYYKVILTVLQNLALI  646
HHV-7      566  L--NFKSQNDFVNMINDLIQCIEEAVSKCISEMRK-TQT-SREQIENCLQSFNIDTTPLSLAFSPFFVFTYYKVILIVLQNLALI  646
HCMV       622  F--SAHGKSDFVAVFSALNKFVDDEALGFVSEVRL---KSSRDEVAGATQAFNLDLNPYAVAFQPLLAYAYFRSVFYVIQNVALI  701
KSHV       553  LRTLIVDIPSFVKVFKGIDSTVEAELLKFINCMIK-NNYNFRENIKSVHHILQFACNVYWQAPCPVFLTLYYKSLLTVIQDICLT  636
EBV        550  E-KEPHGPRDFVKMFKDVDAAVDAEVVQFMNSMAK-NNITYKDLVKSCYHVMQYSCNPFAQPACPIFTQLFYRSLLTILQDISLP  632
Cons_ss:               hhhhhhhhhhhhhhhhhhhhhhhhhhhh       hhh  eeeeeeee         hhhhhhhhhhhhhhhhhhhhh

HSV1       688  QCHGVFAGQSV---EGRNFRNQFQPVLRRRVM-DMFNNGFLSAKTLTVALSEGAAICAPSLTAGQTAPAESSFEGDVARVTLGFP  768
HSV2       688  QCHGVFAGQSV---EGRNFRNQFQPVLRRRVM-DLFNNGFLSAKTLTVALSEGAAICAPSLTAGQTAPAESSFEGDVARVTLGFP  768
VZV        687  QCHCVFYGQQV---EGRNFRNQFQPVLRRRFV-DLFNGGFISTRSITVTLSEG-PVSAPNPTLGQDAPAGRTFDGDLARVSVEVI  766
HCMV       702  TATSYIVDNPLTTNLVSKWMTQHFQSIHGAFSTTSSRKGFLFTKQIKSSKNSDHDRLLDFRLYAQGTYAVVPMEIKLSRLSVPTL  786
HHV-6A     647  VASGHVVDRPCTGNSISKWLVQQYQSLYGTFHSSYLKKGFLNTRTVKVASNVDMEQILDCDLYKSGKYVKTTIQAKLCRLSMQCL  731
HHV-7      647  IGTGYVVDRPCTGNLISKWLMQQYQSLYGAFYNSHFKKGFLNMKTVKIASNVDMEQYIDFNLFKSGKYAKTSIQAKLCRLSMQCL  731
KSHV       637  SCMMYEQDNPAVGIVPSEWLKMHFQTMWTNFKGACFDKGAITGGELKIVHQSMFCDLFDTDAAIGGMFAPARMQVRIARAMLMVP  721
EBV        633  ICMCYENDNPGLGQSPPEWLKGHYQTLCTNFRSLAIDKGVLTAKEAKVVHGEPTCDLPDLDAALQGRVYGRRLPVRMSKVLMLCP  717
Cons_ss:          eeee          hhhhhhhhhhhhhhhhhhhhh       eeeeee   hhh              eeeeeeeeeeee

HSV1       769  KELRVKSRVLFAGASANASEAAKARVASLQSAYQK-PDKRV--DILLGPLGFLLKQFHAAIFPNGKPPGSNQPNPQWFWTALQRN  850
HSV2       769  KELRVKSRVLFAGASANASEAAKARVASLQSAYQK-PDKRV--DILLGPLGFLLKQFHAVIFPNGKPPGSNQPNPQWFWTALQRN  850
VZV        767  RDIRVKNRVVFSGNCTNLSEAARARLVGLASAYQR-QEKRV--DMLHGALGFLLKQFHGLLFPRGMPPNSKSPNPQWFWTLLQRN  848
HCMV       787  IMVRVKNRPIYRAGKG----------NAGSVFFRRDHVPRR--NPAKGCLGFLLYRHHERLFPEC------GLPCLQFWQKVCSN  853
HHV-6A     732  RDFRIKNRPFNKSSKT----------AHNNPYFKKNVKHKK--NPLSGCISFLLFKYHDKLFPNV------KISCLELWQRFLLN  798
HHV-7      732  KDFRVKNRPFNKPNKN----------TQNNPFFKK-VKQKK--NPLSGCLSFLLFKYHERLFPNL------KISCLEFWQRILLN  797
KSHV       722  KTIKIKNRIIFSNS---------TGAESIQAGFMK-PASQRDSYIVGGPYMKFLNALHKTLFPST------KTSALYLWHKIGQT  790
EBV        718  RNIKIKNRVVFTGE-----------NAALQNSFIK-STTRRENYIINGPYMKFLNTYHKTLFPDT------KLSSLYLWHNFSRR  784
Cons_ss:           ee   eee                                     hhhhhhhhhhhhh             hhhhhhhhhh

HSV1       851  QLPARL---LSREDIETIAFIKKFSLDYGAINFINLAPNNVSELAMYYMANQILRYCDHSTYFINTLTAIIAGSRRPPSVQAAAA  932
HSV2       851  QLPARL---LSREDIETIAFIKRFSLDYGAINFINLAPNNVSELAMYYMANQILRYCDHSTYFINTLTAVIAGSRRPPSVQAAAA  932
VZV        849  QMPADK---LTHEEITTIAAVKRFTEEYAAINFINLPPTCIGELAQFYMANLILKYCDHSQYLINTLTSIITGARRPRDPSSVLH  930
HCMV       854  ALPKNVPIGDMGEFNAFVKFLVAVTADYQEHDLLDVAPDCVLSYVESRFHNKFLCYYGF-KDYIGSLHGLTTRLTTQNHAQF-PH  936
HHV-6A     799  NVPKTLDIGNPEEVKTFIKFAFSITNTYDEIDIIDIQPECLSTFIDCYFHNKFLSALGF-HDYLTSLHGLTSKLVTQNPVLF-PV  881
HHV-7      798  NMPKTIDIGNVEDMRSFIKFTFRVTNSYDEIDLLDIQPECLLSFIEYYFHNKLLSVLGY-RDYLTSLHALTSKLVPQNPMLF-PV  880
KSHV       791  TKNPILPGVSGEHLTELCNYVKASSQAFEEINVLDLVPDTLTSYAKIKLNSSILRACGQTQFYATTLSCLSPVTQLVPAEEYPHV  875
EBV        785  RSVPVPSGASAEEYSDLALFVDGGSRAHEESNVIDVVPGNLVTYAKQRLNNAILKACGQTQFYISLIQGLVPRTQSVPARDYPHV  869
Cons_ss:                  hhhhhhhhhhhhhhhhhhhh   ee     hhhhhhhhhhhhhhhh     hh   hhh  e  ee  hhh   h

HSV1       933  WSAQG-----GAGLEAGARALMDAVDAHPGAWTSMFASCNLLRPVMAARPMVVLGLSISKYYGMAGNDRVFQAGNWASLMGGKNA 1012
HSV2       933  WAPQG-----GAGLEAGARALMDSLDAHPGAWTSMFASCNLLRPVMAARPMVVLGLSISKYYGMAGNDRVFQAGNWASLLGGKNA 1012
VZV        931  WIRKDVTS--AADIETQAKALLEKTENLPELWTTAFTSTHLVRAAMNQRPMVVLGISISKYHGAAGNNRVFQAGNWSGLNGGKNV 1013
HCMV       937  VLGASPRFSSPAEFALHVKGLKTAGVPA--PMAATVARESLVRSVFEHRSLVTVPVSVEKYAGINNSKEIYQFGQIGYFSG-NGV 1018
HHV-6A     882  VLDKQPKFSSIQEYLVYVKKLVLDGVPN--PVIASLSKEPNFGTIFTSRSLVTFGLTLEKFVSL-ANREYFQFGQLGWIGG-SGV  962
HHV-7      881  FLKEHPTFSSVQEYVMHVKKLVGNGLKE--PMTASLTKEPNFGSIFTGRSIITFGLMIEKFVSV-ASRDYFHFGQLGWIAG-SGV  961
KSHV       876  LGPVG--LSSPDEYRAKVAGRS--------VTIVQSTLKQAVSTNGRLRPIITVPLVVNKYTGSNGNTNVFHCANLGYFSG-RGV  949
EBV        870  LGTRA--VESAAAYAEATSSLT--------ATTVVCAATDCLSQVCKARPVVTLPVTINKYTGVNGNNQIFQAGNLGYFMG-RGV  943
Cons_ss:        h         hhhhhhhhh  hh         eeee  hhhhhhhh    eeeeeeeeeeee       eeee hhhhh

HSV1      1013  CPLL-------------IFDRTRKFVLACPRAGFVCAASSLGGGAH-ESSLCE-QLRGIISEGGA--AVASSVFVATVK-SLGPR 1079
HSV2      1013  CPLL-------------IFDRTRKFVLACPRAGFVCAASSLGGGAH-EHSLCE-QLRGIIAEGGA--AVASSVFVATVK-SLGPR 1079
VZV       1014  CPLF-------------TFDRTRRFIIACPRGGFICPVTGPSSGNR-ETTLSD-QVRGIIVSGGA--MVQLAIYATVVR-AVGAR 1080
HCMV      1019  ERSLNVS-SM---SGQDYRFMRQRYLLATRLADVLIKRSRRENV-LFDADLIKNRVMLALDAENLDCDPEVMAVYEILS-VR--- 1094
HHV-6A     963  DRNLNPT-SS---ALQDFRFMRQKTIIATKFSEVIVKKVRREAI-MFDTEVVKGKVLSIVENLTNDIDPELLIIAEVMR-DR--- 1038
HHV-7      962  DRNLNPP-SS---GLQDFRFMRQKFVIATKLCDIIVKKVKREAI-VYDVEVIRGKVLNIIESLSNSVNPELLILAEVMK-DR--- 1037
KSHV       950  DRNLRPE-SVPFKKNNVSSMLRKRHVIMTPLVDRLVKRIVGINSGEFEAEAVKRSVQNVLEDRDNP-NLPKTVVLELVKPPRWSS 1032
EBV        944  DRNLLQAPGAGLRKQAGGSSMRKKFVFATPTLGLTVKRRTQAATTYEIENIRA-GLEAIISQKQEE-DCVFDVVCNLVD-AMGEA 1025
Cons_ss:                             eeeeeee      ee           hhhhhhhhhhhhhh       hhhhhhhhhhh h

HSV1      1080  TQQLQIEDWLAL-LEDEYLSEEMMELTARALERGN-GEWSTDAALEVAHEAEALVSQLG---NAGE-----VFNFGDF-----GC 1149
HSV2      1080  TQQLQIEDWLAL-LEDEYLSEEMMEFTTRALERGH-GEWSTDAALEVAHEAEALVSQLG---AAGE-----VFNFGDF-----GD 1149
VZV       1081  AQHMAFDDWLSL-TDDEFLARDLEELHDQIIQTLE-TPWTVEGAL---EAVKILDEKT-TAGDGETPTNLAFNFDSC-----EP 1153
HCMV      1095  EEIPASDDVLFFVDGCEALAASLMDKFAALQE-QGVEDFSLENLRRVLDADAQRLTDA--AGGEVH-----DLSALFAPSGVGAA 1171
HHV-6A    1039  EDKPTMDDMLFFVDGREALAASIMLKLNHLVD-MNVKDFSITNLQSVFETV-----SS--NDAPVY-----DFSEILAEE----- 1105
HHV-7     1038  DSKPTMDDMLFYVDGREPLAKSVMNKIQHLTD-LNVHDFSLSTLLSVFEEQ-----VE--DSAAIY-----DFSELLVEG----- 1104
KSHV      1033  CASLTEEDVIYYLGPYAVLGDEVLSLLSTVGQ--AGVPWTAEGVASVIQDI----IDD--CELQFV-----GPEEPCL-----IQ 1099
EBV       1026  CASLTRDDAEYLLGRFSVLADSVLETLATIAS--SGIEWTAGAARDFLEGV---WGGPGAAQDNFI-----SVAEPVS-----TA 1095
Cons_ss:             hhhhhhhh   hhhhhhhhhhhhhhhh        hhhhhhhhh

HSV1      1150  EDD---NATPF----------GGPG--------APG----PAF---AGRKRAFHGDDPFGE--GPPDKKGDLTLDML 1196
HSV2      1150  ADD---HAASF--------GGLAAA--------AAG----AAG---VARKRAFHGDDPFGE--GPPEKK-DLTLDML 1197
VZV       1154  SHDTTSNVLNI----------SGSN--------ISG----STV---PGLKRPPEDDELFDLS-GIPIKHGNITMEMI 1204
HCMV      1172  SGVG-----------GGGLLLGESVAGNSICFGVPGETGGGCFLVNAGEDEAGGVGGSSGGGGGSGLLPAKRSR--- 1234
HHV-6A    1106  DDQGN-GVLKCDETETETDEPMTKKNRL------------------------------------------------- 1132
HHV-7     1105  NEQGF-GILKCEETEHENEEPSLKKARL------------------------------------------------- 1131
KSHV      1100  GQSVV------------EEL--FPS--------PGVPSLTVGK---KRKIASLLSDLDL------------------ 1133
EBV       1096  SQA---SAGLL--------LGGGGQ--------GSG----GRR---KRRLATVLPGLEV------------------ 1128
Cons_ss:
```

Supplementary Figure S3. **Structure-guided multiple sequence alignment of amino acid sequences of the DNA-binding proteins from representative human herpesviruses.** Alignment was generated in Promals3D[50] using AlphaFold3 models of the proteins. Most conserved residues are highlighted in yellow. Consensus of the secondary structure features among the proteins is given below the alignment, with “h” and “e” representing residues forming α-helices and β-strands, respectively. Viruses included in the alignment: HSV1, herpes simplex virus 1, HSV2, herpes simplex virus 2, VZV, varicella-zoster virus (alphaherpesviruses), HCMV, human cytomegalovirus, HHV-6A, human herpesvirus 6A, HHV-7, human herpesvirus 7 (betaherpesviruses), KSHV, Kaposi’s sarcoma-associated herpesvirus, EBV, Epstein-Barr virus (gammaherpesviruses).

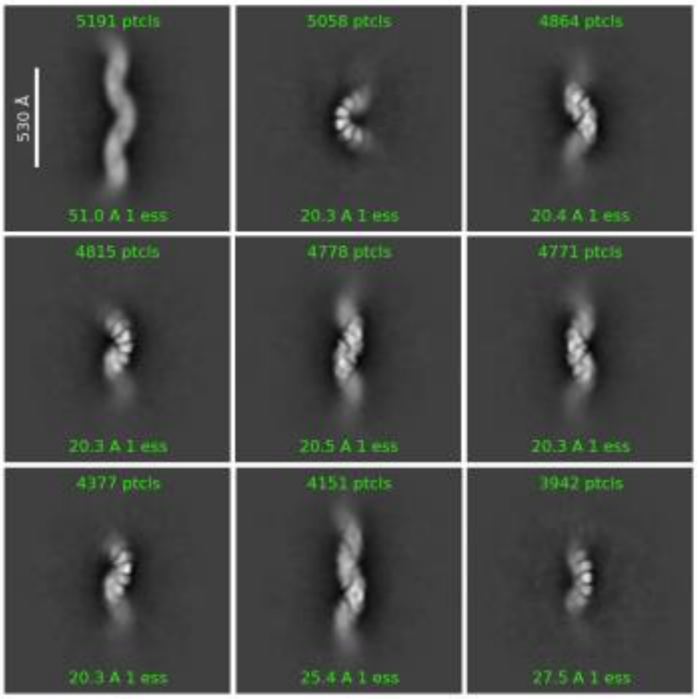


Supplementary Figure S4. **Representative classes from the 2D classification of the BALF2 filament sample analyzed by cryo-EM.**

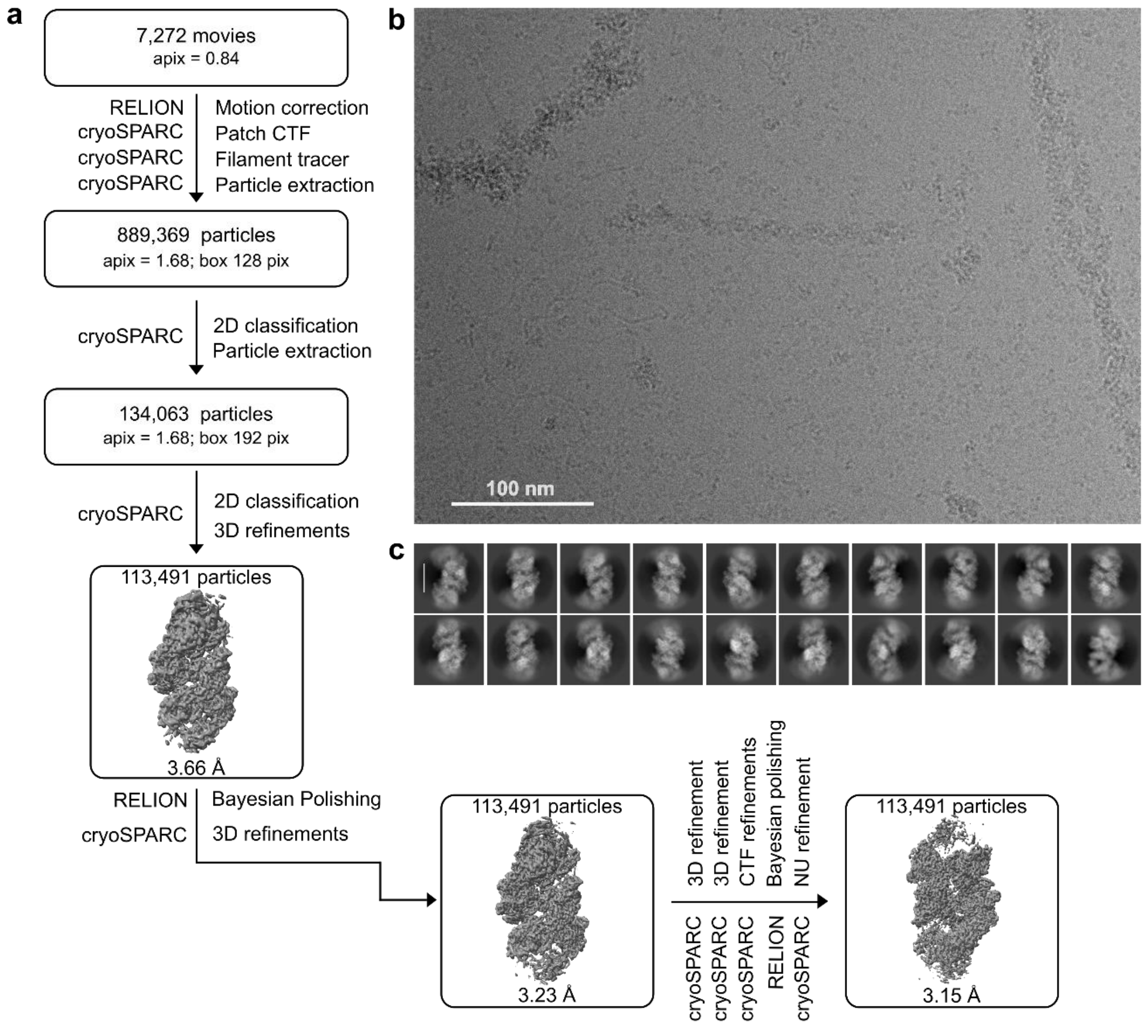


Supplementary Figure S5. **ICP8 Filament cryo-EM data processing. a)** Three-dimensional reconstruction pipeline. **b)** Representative micrograph. **c)** Class averages from the 3rd round of template based 2D classification in cryoSPARC.

Interface 1
(chains A:B)

Interface 2
(chains A:C)

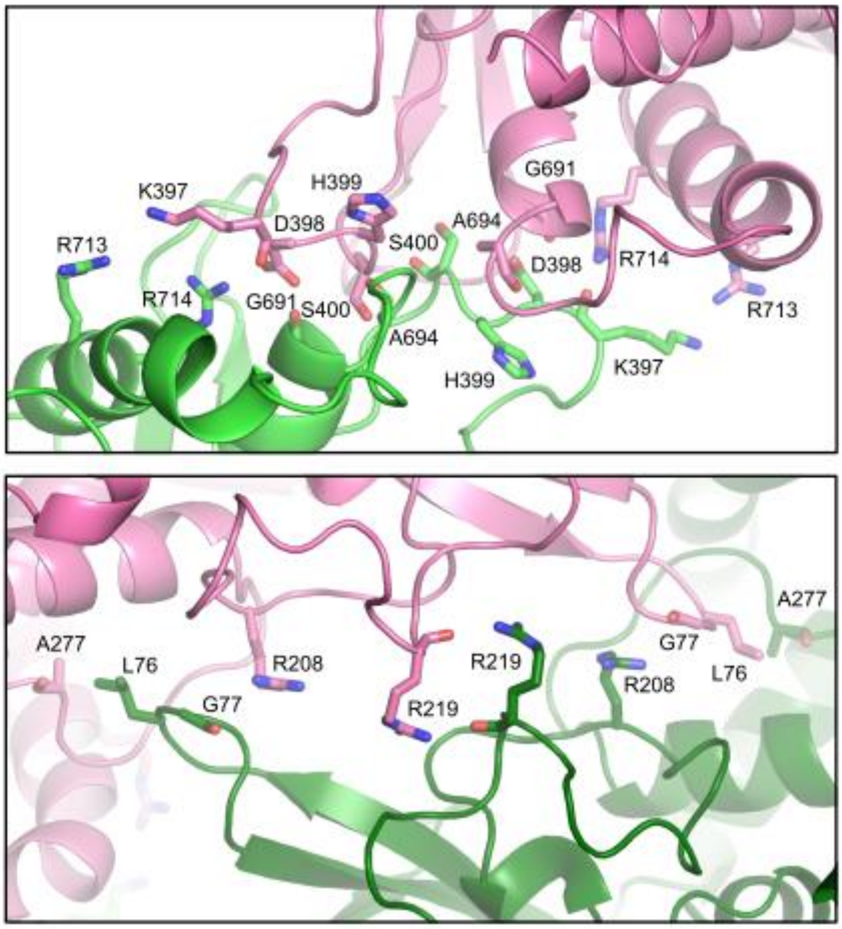


Supplementary Figure S6. **Interfaces between the subunits of the double filament of ICP8.** Protein chains are colored according to the subunit. Residues involved in intersubunit interactions are shown as sticks and labeled.

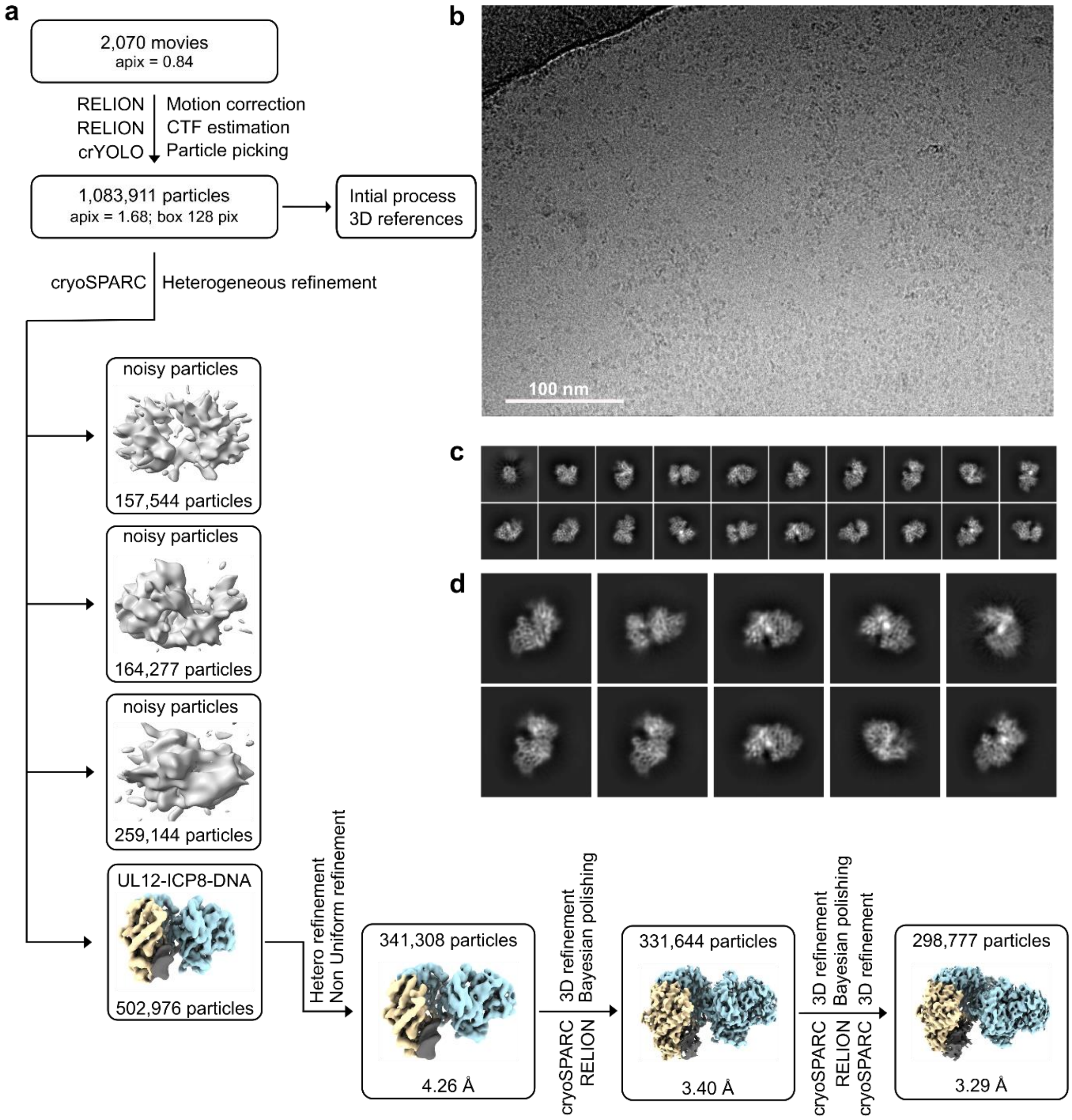


Supplementary Figure S7. **UL12-ICP8-DNA cryo-EM data processing. a)** Three-dimensional reconstruction pipeline. **b)** Representative micrograph. **c)** Class averages from the 3rd round of 2D classification in cryoSPARC. **d)** Enlarged view of class averages from the 3rd round of 2D classification in cryoSPARC.
